# Unraveling the Interplay of Different Traits and Parameters Related to Nitrogen Use Efficiency in Wheat: Insights for Grain Yield Influence

**DOI:** 10.1101/2023.07.03.547462

**Authors:** Gayatri, Puja Mandal, Karnam Venkatesh, Pranab Kumar Mandal

**Affiliations:** ICAR-National Institute for Plant Biotechnology, Pusa Campus, New Delhi-110012, India; Tamil Nadu Agricultural University, Coimbatore-641003, Tamil Nadu, India; ICAR-Indian Institute of Millets Research, Hyderabad-500030, Telangana, India

**Keywords:** Wheat, Nitrogen stress, Nitrogen metabolizing enzymes, Carbon metabolizing enzymes, NUE-related traits

## Abstract

Enhancing Nitrogen Use Efficiency (NUE) in wheat to optimize grain yield is a significant challenge. To address this challenge, a comprehensive study was conducted to investigate various morphological, biochemical, molecular parameters, and agronomic traits related to NUE. By examining various traits under both optimum-N (ON) and stressed-N (SN) conditions, the study explores the interrelationships among these traits, providing novel insights not previously reported. A set of 278 diverse wheat genotypes were assessed, encompassing eight NUE-related traits: Grain Yield, Biomass, Grain nitrogen, N at head, N at harvest, N-uptake, Nitrogen Uptake Efficiency, Nitrogen Utilization Efficiency, and NUE under both ON and SN conditions in the field. The findings demonstrated a significant positive correlation between grain yield and all NUE-related traits, highlighting their significance in comprehending the biological NUE of wheat plants. Notably, the study identified N-uptake and N-uptake related traits as key factors influencing the impact of soil nitrogen status on yield and associated parameters. These traits hold particular importance for selecting wheat genotypes with optimal yield and NUE in wheat cultivation. To complement the field data, representative genotypes were further subjected to a hydroponics experiment under absolute nitrogen control. This experiment provided insights into the effects of nitrogen stress on morphological parameters and the performance of eight essential nitrogen and carbon metabolizing enzymes. Correlation analysis highlighted the substantial influence of four key N-metabolizing enzymes, namely Nitrate Reductase, Glutamine Synthetase, Glutamate Oxo-Glutarate Amino Transferase, and Glutamate Dehydrogenase, on grain yield. Additionally, this study underscored the direct and indirect associations between seedling parameters and field traits, emphasizing the significance of shoot and root length parameters in nitrogen acquisition under nitrogen stress. In conclusion, these findings offer valuable insights into the intricate network of traits and parameters that influence wheat grain yield under varying nitrogen regimes. This knowledge can aid in the selection of wheat genotypes with enhanced NUE and grain yield, particularly in scenarios of reduced nitrogen application.

**Key Points:**

- A comprehensive study in field and hydroponics conditions revealed differential responses of various morphological, biochemical, molecular parameters, and agronomic traits to different nitrogen levels.
- N-uptake related traits in field condition and chlorophyll content and morphological parameters in hydroponics condition were found as essential factors contributing to variations under both optimum and stressed N conditions.
- Among the parameters observed in the seedling stage, SL and RL, along with the enzymes NR, GS, GOGAT, and GDH, demonstrated their influence on GY.

## Introduction

Over the past 50 years, there has been a strong correlation between the enormous intake of nitrogen (N) fertilizers and the rise in global agricultural productivity. However, Nitrogen use efficiency (NUE) in many cereal crops is less than 50% (Raun & Johnson, 1999; F. ZHANG et al., 2015; H. ZHANG et al., 2018). Therefore, a high amount of nitrogen fertilizers was used to increase crop productivity, which in turn has an adverse impact on the environment as well as human health (Qiao et al., 2012; Sieling & Kage, 2008; H. ZHANG et al., 2018). Therefore, enhancement of NUE is a major concern for maintaining sustainable yield using less application of N fertilizers. Moreover, NUE is a complex quantitative trait that is influenced by biological, physiological, environmental, genetic, agronomic, and developmental factors (Congreves et al., 2021; Hawkesford, 2017; Hawkesford & Griffiths, 2019; Hirel et al., 2011). Hence, the improvement of NUE in plants has multidimensional dependency and interrelations. Perhaps understanding these factors and their interrelations and correlations could pave the way towards improving NUE because none of the single factors can address all the concerns of NUE. Several approaches have been used to study the impact/response of different factors contributing to NUE, e.g., different fertilizers (Bargaz et al., 2018; Li et al., 2018; Quevedo-Amaya et al., 2020) and crop management strategies (Cui et al., 2018; Guo et al., 2017; Santiago-Arenas et al., 2020), transgenic approaches (Mandal et al., 2018; Raghuram et al., 2019; Sinha et al., 2020), functional genomics (Pathak et al., 2020), genome-wide association mapping (Tang et al., 2019), evaluation of phenotypic and physiological parameters (Chardon et al., 2010; Jain et al., 2011; Mukami et al., 2019), etc. These strategies and approaches were mostly attempted with a few traits/parameters without addressing the various aspects related to NUE. Moreover, it is still unknown which traits/parameters are major contributors to wheat NUE. Additionally, plant NUE is inherently a complex trait because of having multiple physiological processes, including N uptake, assimilation, translocation, and remobilization governed by several interacting genetic and environmental factors (Giambalvo et al., 2010). Hence, to enhance NUE, it is imperative that a more profound knowledge of these processes and the underlying mechanisms governing them be fully understood (Kant et al., 2011).

For most plants, nitrate is the major N source which is uptaken by the roots using different nitrate transporters. The uptaken nitrate is then further processed through the N-assimilation pathway using different key enzymes, i.e. nitrate reductase (NR), nitrite reductase (NiR), glutamine synthetase (GS), glutamate synthase/ glutamate oxo-glutarate aminotransferase (GOGAT), glutamate dehydrogenase (GDH) (Foulkes et al., 2009; Jain et al., 2011). NR is the first enzyme involved in N assimilation, which reduces the nitrate into nitrite (Crawford & Arst Jr, 1993; Lea, 1993; Masclaux-Daubresse et al., 2010). Nitrite is further reduced to ammonium by the action of NiR, and ammonia is assimilated into organic form as glutamine and glutamate in chloroplast through the glutamine synthetase⁄glutamate synthase (GS⁄GOGAT) cycle (Foyer et al., 2011; Masclaux-Daubresse et al., 2010; Suzuki & Knaff, 2005), which provides nitrogen for the biosynthesis of amino acids, nucleic acids, and other nitrogen containing compounds such as chlorophyll. The controversy over the importance of glutamate dehydrogenase (GDH) enzyme still exists for ammonia assimilation or carbon recycling in plants (Dubois et al., 2003; Tercé-Laforgue et al., 2004). GDH involves an alternative pathway for N assimilation (Hirel et al., 2011; Shrawat & Good, 2008). GDH catalyzes the amination of 2-oxoglutarate into glutamate (anabolic reaction) and/or the deamination of glutamate into ammonia and 2-oxoglutarate (catabolic reaction) (Dubois et al., 2003; Lea et al., 1990). Numerous investigations have shown that C and N metabolism is tightly regulated (Scheible et al., 2000; Zheng, 2009), and this was also reported that the regulation of GS, GOGAT, PK, CS, and ICDH are controlled by a common transcription factor *Dof1* (Bharati et al., 2022; Kumar et al., 2009; Yanagisawa, 2002, 2004) (Figure 1). Hence, studying these enzymes involved in C and N metabolism under the same level of N and in the same genetic background may provide a better understanding. During nitrate assimilation, carbohydrate synthesis is decreased, and more carbon is converted via glycolysis to phosphoenolpyruvate (PEP) by the enzyme phosphoenol pyruvate carboxylase (PEPC) and enters organic acid metabolism (Scheible et al., 2000; Stitt & Krapp, 1999). Organic acid metabolism is involved in N-assimilation where PEPC operates together with pyruvate kinase (PK), the mitochondrial citrate synthase (CS), pyruvate dehydrogenase, and the cytosolic NADP-dependent isocitrate dehydrogenase (NADP-ICDH) (Scheible et al., 1997; Stitt & Krapp, 1999) to provide 2-oxoglutarate, which is the primary carbon acceptor for ammonium. Incorporation of inorganic N into organic compounds in plants requires the C skeleton derived from the mitochondrial respiratory system as well as organic acid metabolism. Nitrate supply has been shown to result in the decrease of starch synthesis and diversion of C towards the conversion of organic acids into amino acids (Scheible et al., 2000).

**Figure 1.**
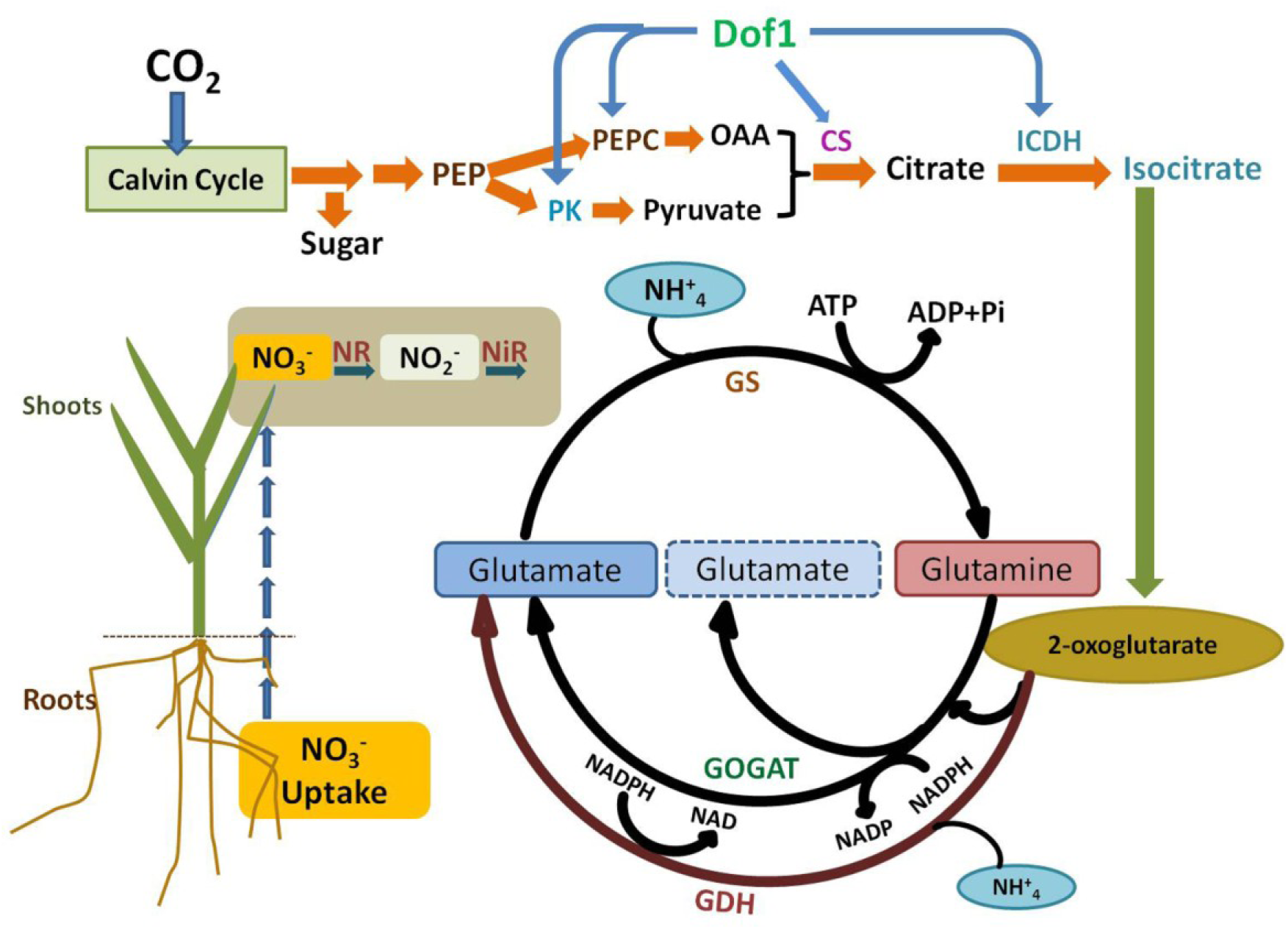
Simplified whole plant view of tightly regulated C and N metabolism.

Hence, along with the yield and morphological parameters, it is important to study the key enzymes involved in the C/N assimilation pathway for a better understanding of the NUE (Grain yield per unit of N) of the crop. Wheat is the most important cereal crop in the world, and NUE is a critically important concept for growing the crop under judicious N. Several studies have investigated the differential N responses of various key enzymes in wheat genotypes but are limited to a few genotypes only (Chandna et al., 2010; Jain et al., 2011; Kaur et al., 2015). Thus, the characterization of these vital enzymes involved in C/N assimilation is important in relation to NUE. None of the reported studies have comprehensively evaluated NUE-related parameters, including these enzymes involved in C/N assimilation, neither in wheat nor in any other crop. Further, the correlations between different field traits with the parameters derived from the experiment under the controlled condition are important in identifying and improving NUE in response to low and optimum N conditions. Hence, to know the contribution of each trait/parameter under N-optimum and N-stress conditions involving a set of diverse genotypes, we have studied important N-responsive parameters of wheat both under field as well as laboratory/controlled conditions.

NUE is a term commonly used by researchers to describe how effectively plants utilize nitrogen from the soil. There are several definitions of NUE that have been proposed in the literature (Congreves et al., 2021; Dobermann, 2007; Ernst et al., 2020; Fageria et al., 2008; Good et al., 2004; Ladha et al., 2005), each with their own advantages and limitations. We felt that grain yield per unit of available nitrogen is the most appropriate with respect to agriculture. However, the focus is neither an increase in yield with the additional amount of nitrogen nor a significant decrease in yield with very low available nitrogen. In today’s context, the ideal situation would be the reduction of N-fertilizer application without compromising yield. Keeping this in mind, we started our field experiment with 278 wheat genotypes and later selected 17 diverse genotypes and investigated various N-responsive parameters under N-optimum and N-stressed conditions, which are likely to determine the yield or NUE of wheat. Plant biomass, grain yield, N content, Nitrogen uptake, Nitrogen-uptake Efficiency (NUpE), Nitrogen Utilization Efficiency (NUtE), and NUE were studied at the full maturity stage in field condition, while plant biomass, chlorophyll pigments were studied in hydroponics conditions. Although the *in vitro* assays of a few enzymes and their transcript expression was reported in a few wheat genotypes (Chandna et al., 2010; Jain et al., 2011; Kaur et al., 2015; Sinha et al., 2015), we have assayed eight enzymes, those are related to N and C metabolism, and also studied their gene expression. Probably this would be the first report where these many parameters from field and hydroponics experiments have been included to find the major contributor towards wheat N use or yield.

## Materials and Methods

### Plant materials

A diverse collection of 278 bread wheat (*Triticum aestivum* L.) genotypes (for simplicity were numbered as No. 1–278 and provided the genotype names in Supp Table 1 were used for the field experiment.

For laboratory experiment, 17 genotypes that represented a subset of the above genetic material (Supp Table 1) were used, and other details, including their pedigree, have been provided in Supp Table 2.

### Experiments Design

#### Field experiment and measured traits

For the field experiment, 278 genotypes (Supp Table 1) were grown at ICAR-Indian Institute of Wheat & Barley Research (IIWBR), Karnal, India (29.7029° N, 76.9919° E) in 2013–14. These genotypes were grown in plots (1m x 0.8 m) under a standard pack of practices at two levels of nitrogen with three replications each (Bharati et al., 2022). One set of the genotypes was grown in normal plots (having approximately 150 kg/ha of available soil nitrogen) with 150kg/ha nitrogen through fertilizer application, and the other set was grown in N-stressed plots (having 100 kg/ha of soil available nitrogen) with no additional nitrogenous fertilizer. The N-stress plots were established by ceasing all forms of external nitrogen supply since 2012, and *Sorghum bicolor* was grown in these plots to speed up the depletion of nitrogen (Sevanthi et al., 2021) during the *Kharif* season.

At the time of harvest, the above-ground part of the plants excluding the grains, was considered as the biomass (BM). After threshing and separating the grains from the panicles, Grain Yield (GY) was measured. BM, GY, as well as other parameters were calculated per plot basis. Both available soil and plant nitrogen (Straw and Grain N) was estimated by using the Micro-Kjeldahl method (Kjeldahl digestion by KMnO_4_ followed by distillation and titration). The following derived parameters were determined (Hawkesford & Griffiths, 2019; Moll et al., 1982) using these formulae:

at head = N%Head * BM/100
at harvest = N% harvest * BM/100
N-uptake = Grain N (GN) + N at Harvest
= N uptake/ (Soil N+ Applied N)
= (GY)/N uptake
= GY/ (available soil N content + N (fertilizer) applied) or NUE = NUpE x NUtE

#### Hydroponics experiment

Representative 17 genotypes were grown hydroponically for 15 days under optimum N (ON: 8 mmol N /L) and Stressed N (SN: 0.08 mmol N /L) conditions in the culture room (Gayatri et al. 2019, 2021). Seedlings were harvested after 15 days of growth, and shoots and roots were separated and stored at −80°C after snap freezing in liquid nitrogen. Experiments were conducted in a completely randomized design with three replications. We recorded the data from three biological replicates for all the parameters, and each biological replicate consisted of a pool of three seedlings.

#### Biomass

Separated shoot and root samples were used for the measurement of shoot length (SL), shoot fresh weight (SFW), shoot dry weight (SDW), root length (RL), root fresh weight (RFW), and root dry weight (RDW) using the standard protocols (Sinha et al., 2015).

#### Chlorophyll and carotenoid contents estimation

Chlorophyll and carotenoid contents were measured from the fresh leaf tissues (Hiscox & Israelstam, 1979).

#### *In Vitro* assays of key enzymes involved in C/N metabolism

Eight key enzymes involved in C and N metabolism, i.e., NR, NiR, GS, GOGAT, GDH, CS, ICDH, and PK were assayed in the freshly harvested shoot tissues from 15 days old wheat seedlings. NR, GS, GOGAT, GDH, and PK enzymes were assayed using the protocol mentioned by Sinha et al., 2015. NiR, CS, and ICDH enzymatic assays were carried out by using the methods of Grover and Lohda (1998), Srere et al. (1963), and Bergmeyer and Berm (1974), respectively. Total soluble protein was estimated by using the method of Lowry et al. (1951). Enzyme activity calculated here for each enzyme is activity/mg of protein, i.e, specific activity.

### Gene expression study of the enzymes

#### Total RNA Isolation and cDNA Synthesis

Total RNA extraction was done using the NucleoSpin™ RNA Plant Kit from the stored shoot tissues, and the cDNA was synthesized using SuperScript®III First-strand Synthesis SuperMix (Invitrogen™, USA) kit using oligodT primer (Gayatri et al., 2019).

#### Quantitative Real-Time PCR (qRT-PCR)

The expression pattern for all genes was determined using qRT-PCR (Gayatri et al., 2021). RT-qPCR primers for all the genes were designed from the conserved region of the coding sequences of a gene (Supp Table 3). The expression of each gene was calculated as relative fold change using the relative 2^−ΔΔCt^ method (Gayatri et al., 2021; Schmittgen & Livak, 2008).

### Statistical analysis

The data obtained were analyzed statistically by analysis of variance (ANOVA) for each parameter separately using the MSTATC program. The significance of differences among genotypes will be determined by the least significance difference (LSD) calculated at P<0.05. In the case of the enzymatic assay, factorial analysis was carried out (CRD, 2 factor analysis with treatment and genotypes as two factors) to obtain ANOVA. Subsequently, the Least Significant difference at 5% was calculated for treatment x genotype combination, and the range was derived through a range test, and the value was expressed in alphabets. Two sample t-test was employed to compare the means of genotypes at ON and SN conditions, with the significance level set at P=0.05. Pearson’s correlations among the traits were performed by using the metan package (Olivoto & Lucio, 2020), principal component analysis (PCA) was performed by using the factoextra package based on the ggplot2 package (Wickham, 2009), and stepwise regression analysis was done using MASS package in R software v.3.4.3.

## Results

### Field Experiment

The mean values of field traits ranged from 243.1-705.8 g/m^2^ for GY, 656.8-1825 g/m^2^ for BM, 1.2-16.0 g/m^2^ for GN, and 8.8-37.6 g/m^2^ for N at Head, 2.4-16.3 g/m^2^ for N at Harvest (straw), 3.6-26.3 g/m^2^ for N uptake, 11.9-78.3 for NUpE, 15.0-89.5 for NUtE, 8.1-23.5 for NUE under ON condition (Supp Table 4). Similarly, the mean values of morphological parameters under SN ranged from 74.0-276.7 g/m^2^ for GY, 200.3-857.7 g/m^2^ for BM, 1.2-5.5 g/m^2^ for GN, and 2.3-10.1 g/m^2^ for N at Head, 0.5-2.8 g/m^2^ for N at Harvest, 1.7-6.8 g/m^2^ for N uptake, 17.2-67.5 for NUpE, 26.8-83.8 for NUtE, 7.4-27.7 for NUE (Supp Table 4).

### Significance and Effect of N-stress on NUE-related traits measured at field conditions

Based on the field data of 278 diverse genotypes grown under N-optimum and N-stress conditions, we have summarized the variability of 9 N-stress responsive traits in the form of a box plot (Figure 2) (Supp Table 4). The majority of these traits showed significant differences between ON and SN conditions. T-test revealed that the values of most of these traits under N-stress were significantly (P=0.05) reduced, except NUtE and NUE (Figure 3). The mean value of all the traits was calculated, and it was found that the value of the majority of the traits has reduced under SN (reduction of GY 59.7%, BM 59.2%, GN 64.8%, N at Head 77.5%, N at Harvest 76%, N-uptake 69.2%, NUpE 7.5%) (Figure 3). However, the two derived parameters NUtE and NUE values were projected to be higher (NUtE 20.6%, NUE 20.8%) due to low available nitrogen and low uptake (which is there in the denominator of the formulae).

**Figure 2:**
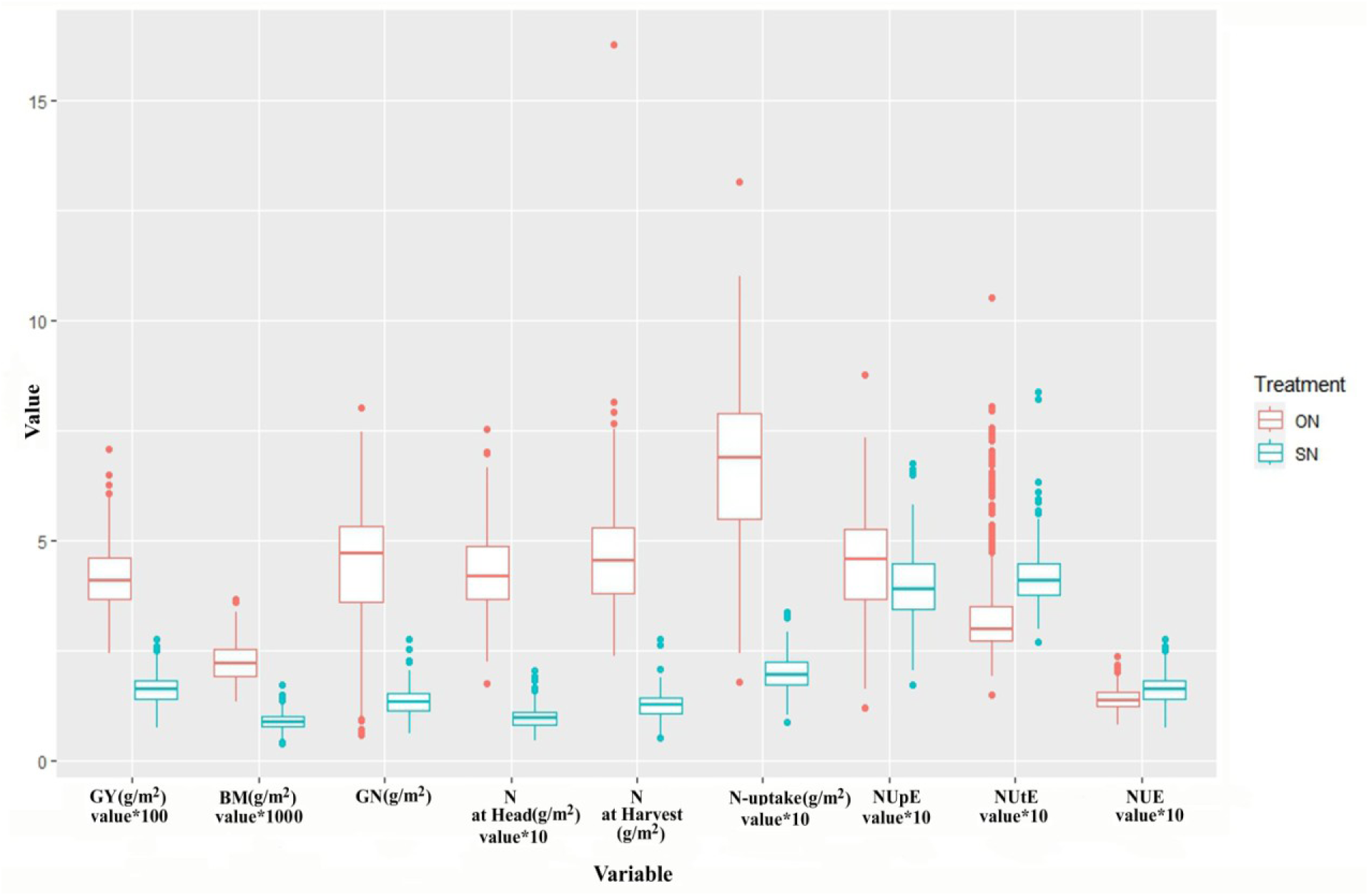
Summary of data observed for 9 NUE traits under ON (optimum-N) and SN (stressed-N) conditions in the field. Boxplots of 9 NUE traits showing variation within 278 selected wheat genotypes. Keys: GY: grain yield; BM: biomass, GN: grain nitrogen, NUpE: Nitrogen Uptake Efficiency; NUtE: Nitrogen Utilization Efficiency; NUE: Nitrogen Use Efficiency

**Figure 3:**
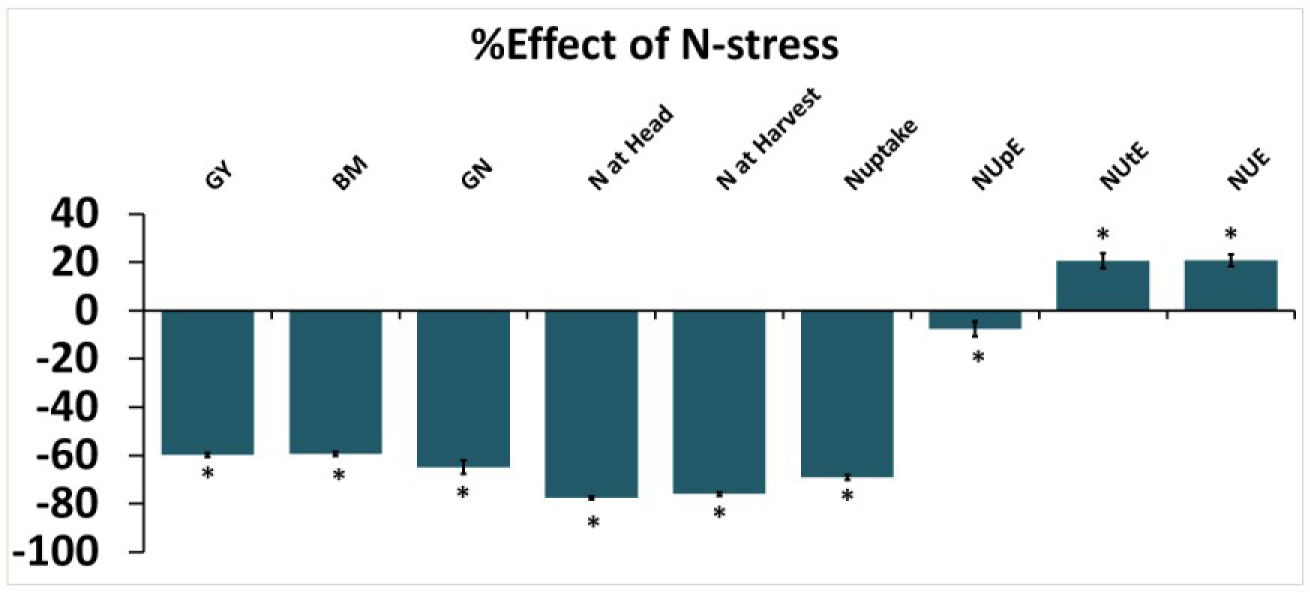
Differential effect of N-stress on various NUE traits. The % effect of stressed-N (SN) minus optimum-N (ON) on each of the 9 traits were calculated using the mean values of 834 replicates and pooled for all the genotypes. * represents the significance at P < 0.05. Keys: GY: grain yield; BM: biomass, GN: grain nitrogen, NUpE: Nitrogen Uptake Efficiency; NUtE: Nitrogen Utilization Efficiency; NUE: Nitrogen Use Efficiency

### Principal Component Analysis (PCA) for field traits

PCA was used to separate the individual contribution of each trait to the total phenotypic variation noticed among 278 genotypes. In our study, PCA was carried out to determine the traits responsible for different degrees of phenotypic variation among the genotypes. The PCA grouped the estimated wheat field traits/variables into two main components under ON and SN conditions. Under ON condition, PC1 accounted for about 52.4% of the variation; PC2 for 30.5% (Supp File 1: Supp Table 5). While under SN condition, PC1 accounted for about 68% of the variation; PC2 for 18.1% (Supp File 1: Supp Table 6). Under both ON and SN conditions, all traits, except NUtE, contributed positively to PC1 (Figure 4). The major contributors for PC-1 were N-uptake and NUpE (acquisition) under both conditions (Figure 4). Based on the PCA, under the ON condition, N at head and N at harvest showed a strong correlation with GY, NUE, and BM but not with GN (Figure 4a), while under the SN condition, N at head and N at harvest showed a strong positive correlation with N uptake, NUpE, GN, BM rather than with GY and NUE (Figure 4b). Under both conditions, NUtE showed a negative correlation with NUpE and GN.

**Figure 4:**
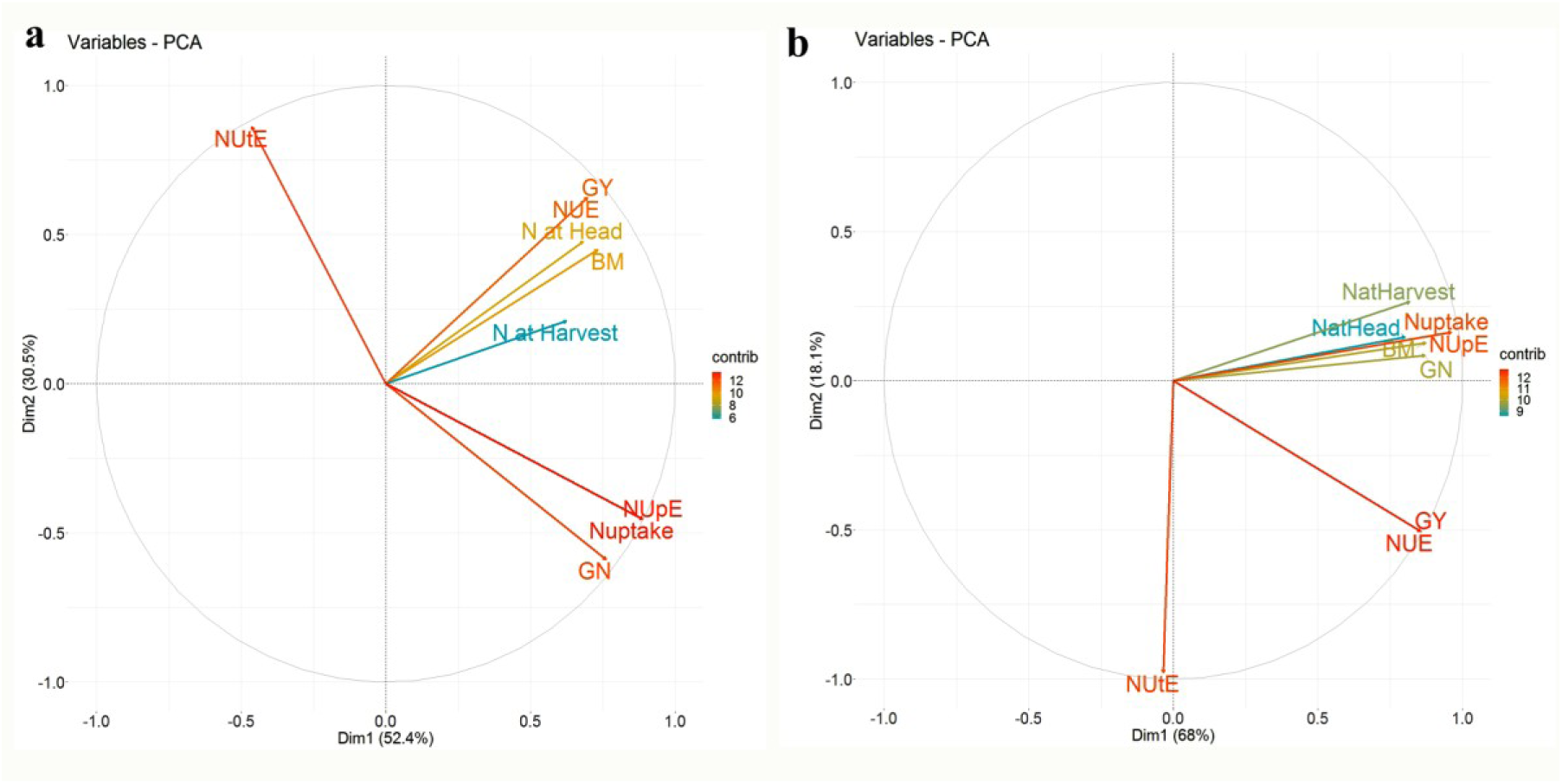
Biplot for NUE traits measured under field (a) ON and (b) SN conditions. Keys: GY: grain yield; BM: biomass, GN: grain nitrogen, NUpE: Nitrogen Uptake Efficiency; NUtE: Nitrogen Utilization Efficiency; NUE: Nitrogen Use Efficiency

### Correlation between different traits measured at field conditions

The results obtained for correlation among 9 different traits under ON and SN are presented in Figure 5. NUE was calculated as GY divided by available soil N (which is a constant value across the genotypes); therefore, the correlation analysis of NUE and GY with other traits was the same, having the same r^2^ value (Figure 5). Likewise, NUpE has been derived from N-uptake; hence these two traits showed the same r^2^ value when correlation was derived with other parameters.

**Figure 5:**
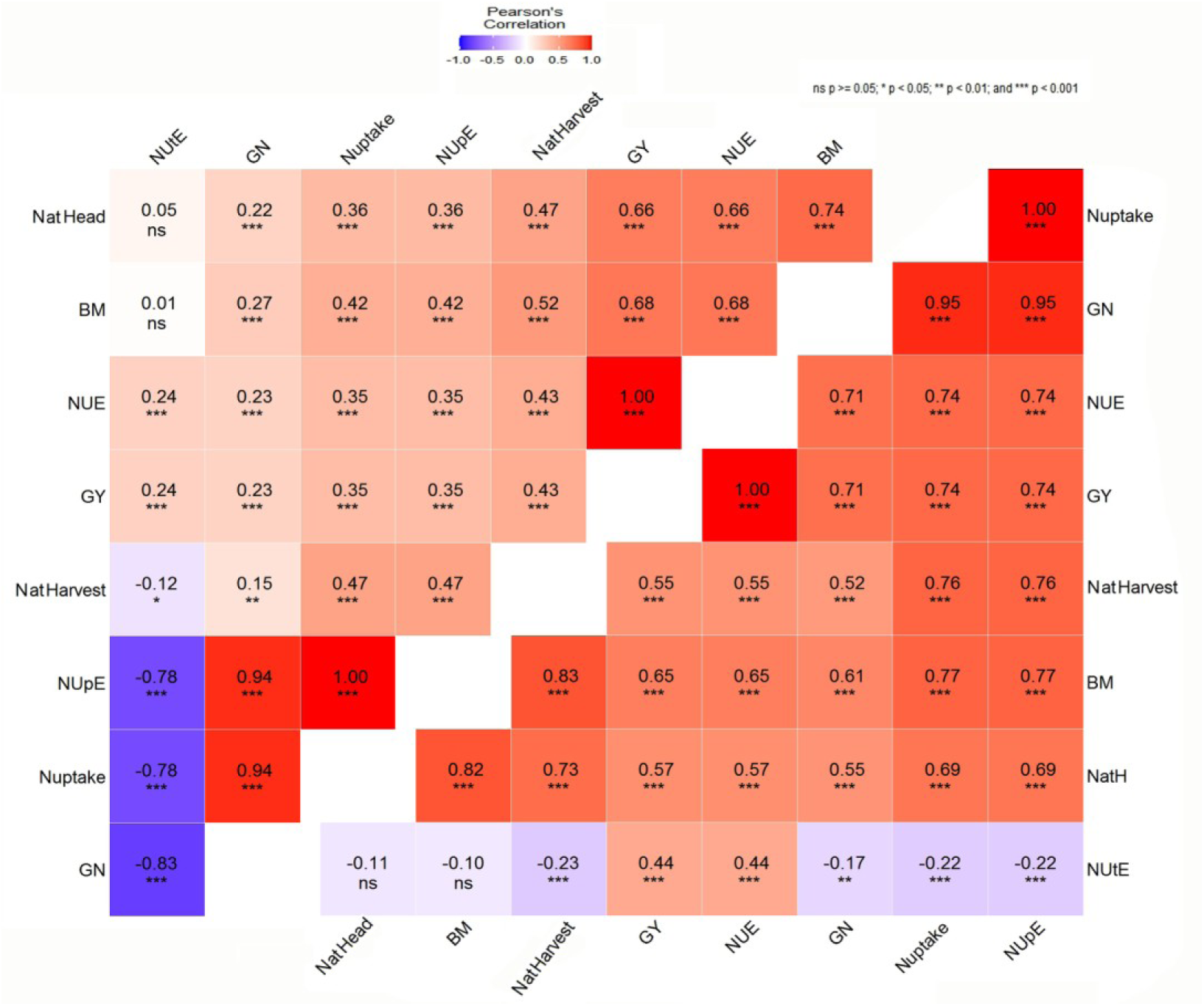
Correlation analysis among the 9 NUE traits at the field condition. The upper triangle shows correlation analysis at the ON condition, and the lower triangle shows correlation analysis at the SN condition. The red color indicates a strong positive correlation, while the blue color indicates a strong negative correlation. Keys: GY: grain yield; BM: biomass, GN: grain nitrogen, NUpE: Nitrogen-uptake efficiency; NUtE: Nitrogen Utilization Efficiency; NUE: Nitrogen Use Efficiency

Under both ON and SN conditions, all traits showed correlation with each other, except NUtE was not correlated with N at Head and BM (Figure 5). NUtE showed a strong negative correlation with GN.

### Hydroponics Experiment

17 genotypes that represented a subset of the above genetic material were picked based on their diversity of GY (Supp File 2: Supp Figure 1) and BM (Supp File 2: Supp Figure 2) under both N-optimum and N-stressed conditions. Based on the hydroponics data of 17 selected genotypes (Supp File 3: Supp Table 7) grown under N-optimum and N-stress conditions, we have summarized the variability of 19 parameters (11 morpho-physiological parameters and 8 enzyme assays) in the form of a box plot (Figure 6).

**Figure 6:**
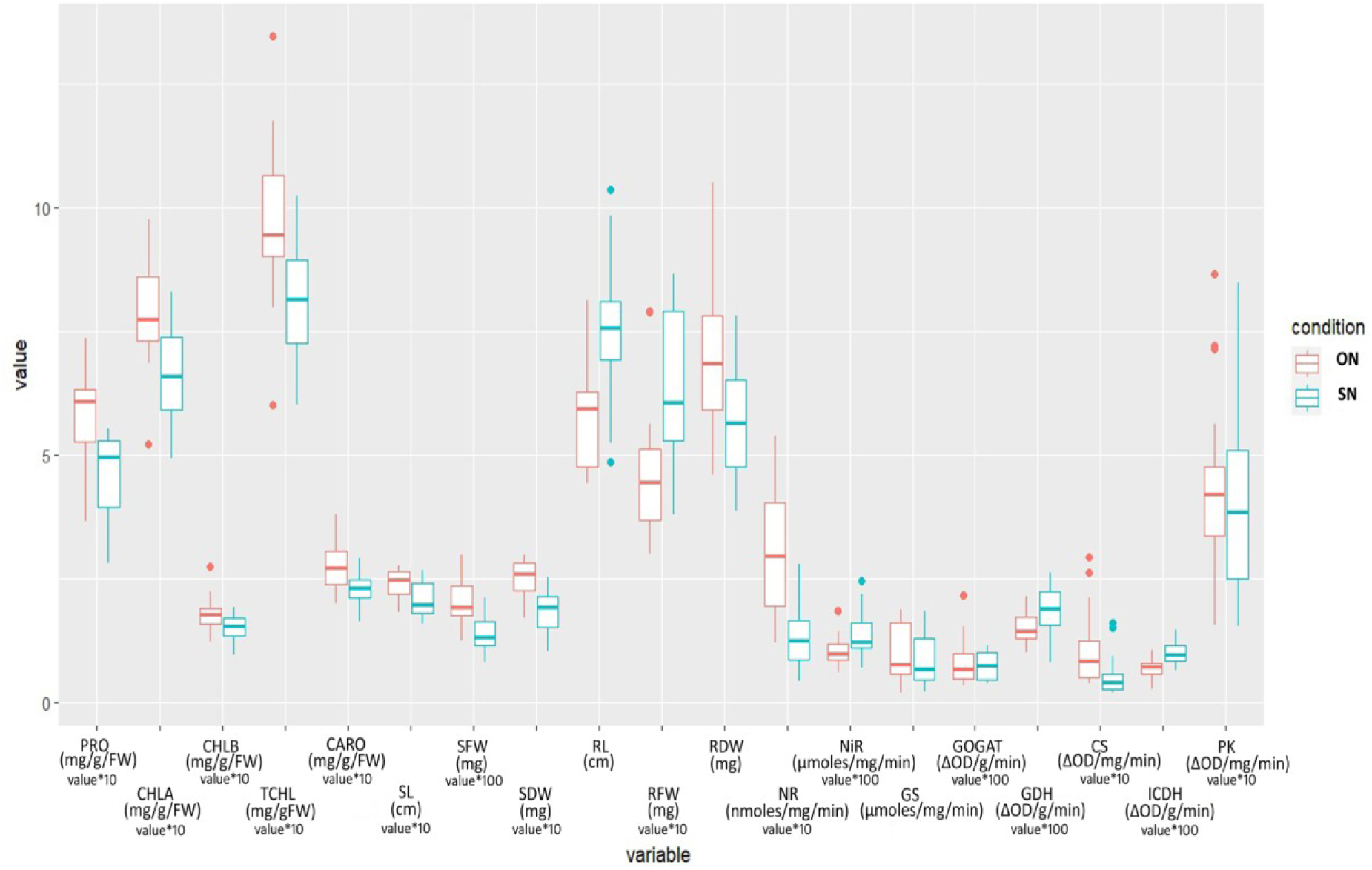
Summary of data observed for morphological parameters and enzyme activities under ON (optimum-N) and SN (stressed-N) conditions in hydroponics. Keys: PRO: protein; CHLA: Chlorophyll A; CHLB: Chlorophyll B; TCHL: Total Chlorophyll; Caro: Carotenoids; SL: shoot length; SFW: shoot fresh weight, SDW: shoot dry weight, RL: root length; RFW: root fresh weight

### Morphological parameters

The mean values of morphological parameters ranged from 18.23-27.6 cm for SL, 123.8-297.6 mg for SFW, 17-29.73 mg for SDW, 4.5-8.13 cm for RL, 30.03-79.06 mg for RFW, and 4.59-10.50 mg for RDW under ON condition (Supp File 3: Supp Table 7). Similarly, the mean values of morphological parameters under SN ranged from 15.77-26.7 cm for SL, 81.07-212.1 mg for SFW, 10.33-28.3 mg for SDW, 4.86-10.37 cm for RL, 38.08-86.56 mg for RFW, 3.86-7.81 mg for RDW (Supp File 3: Supp Table 7). The result shows the variability of the genotypes for all parameters.

### Photosynthesis Pigments

In the ON conditions, Car, Chla, Chlb, and ChlT ranged from 0.19–0.38, 0.52–0.97, 0.12–0.27, 0.6–1.34 mg g^-1^FW respectively; in contrast, under SN conditions, mean values of Car, Chla, Chlb, and ChlT ranged from 0.16–0.28; 0.49–0.83; 0.09–0.19 and 0.60–1.02 mg g^-1^FW respectively (Supp File 3: Supp Table 7).

### Total soluble protein

Soluble protein content was found in a range of 36.5-73.47 mg g^-1^FW, and 28.06-55.3 mg g^-1^FW (Supp File 3: Supp Table 7) under ON and SN conditions, respectively.

### Activities of N and C-assimilation enzymes

In ON conditions, NR activity was found in the range of 11.9-53.9 nmoles nitrate reduced/mg protein/min. While under N stress conditions, the range of NR activity was from 4.2-27.8 nmoles nitrate reduced/mg protein/min (Supp File 3: Supp Table 7). But in the case of NiR enzyme, all genotypes showed a contrasting behavior, i.e., under SN; NiR activity was found to be high. NiR activity was found in a range of 58.89-185.7 and 69.12-245.14 μmoles nitrite reduced/mg protein/min under ON and SN conditions, respectively.

The GS, GOGAT, and GDH activities did not show any regular pattern among all genotypes. All the genotypes showed GS activity in a range of 0.19-2.08 and 0.2-1.8 μmoles γ-glutamyl hydroxamate formed/mg protein/min (Supp File 3: Supp Table 7), GOGAT activity in a range of 3.26-21.57 and 3.74-11.45 change in OD/min/g protein (Supp File 3: Supp Table 7) and GDH activity in a range of 10-21.33 and 8.33-26 change in OD/min/g protein (Supp File 3: Supp Table 7) under ON and SN respectively. PK specific activity did not show any regular pattern among all genotypes. PK activity was found in a range of 0.16-0.86 and 0.15-0.85 change in OD/min/mg protein under ON and SN conditions, respectively (Supp File 3: Supp Table 7). The range of activity of CS was 0.038-0.293 and 0.018-0.16 change in OD/min/mg protein under ON and SN conditions, respectively. The range of activity of ICDH were 2.61-10.45 and 6.49-14.68 change in OD/min/g protein under ON and SN condition, respectively (Supp File 3: Supp Table 7). Logically, when N-is less, none of the enzymes involved in N-assimilation should have higher activity, but NiR and GDH, and ICDH are the exceptions.

### Expression of N and C-assimilation genes

The result shows that NR and NiR both genes were downregulated in all genotypes (Figure 7). Whereas *GS, GOGAT, GDH, CS*, *ICDH*, *and PK* gene expression was found genotype dependent, which showed up and down-regulation among these diverse genotypes (Figure7).

**Figure 7:**
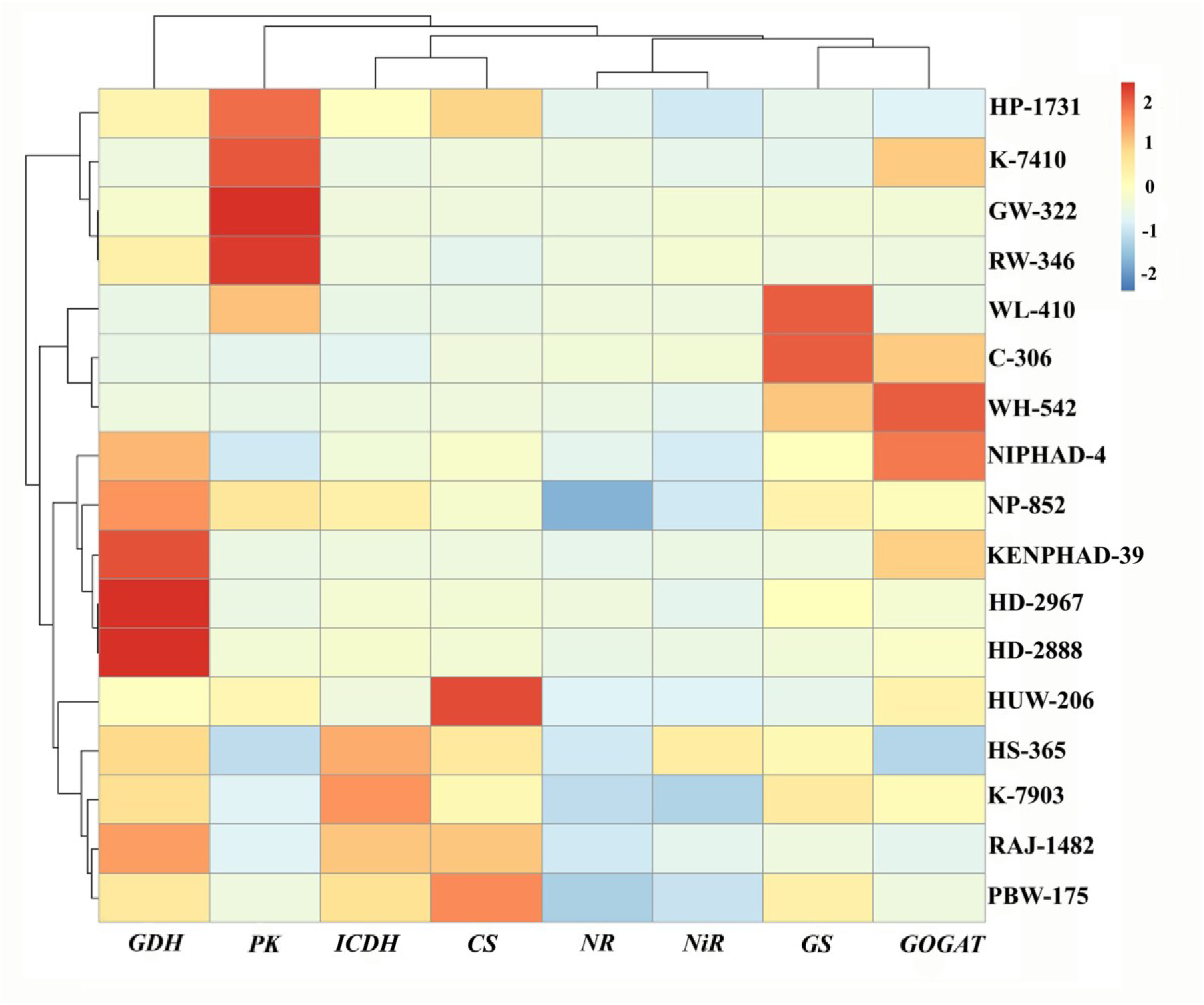
Heat map showing differential expression patterns of eight genes involved in N and C metabolism in seedlings’ shoots between ON and SN conditions. The color bar of the heat map shows the fold change; thereby, red color represents upregulation, white represents no change, and blue color represents downregulation.

### Magnitude and Significance of N-stress on different parameters

a t-test revealed that the effect of N-stress was significant (P=0.05) in all the parameters except GS, GOGAT, and PK enzyme activities as well as their transcript level (Figure 8) measured hydroponically. For this purpose, mean data values for each of the parameters were averaged for all the genotypes, and the % effect of N on each parameter (SN minus ON) was plotted (Figure 7). It is evident that N-stress affects different parameters to different extents. The % effect of N-stress was negative on most of the parameters by -0.84 to -79.8% except NiR_act, GDH_act, ICDH_act, RL, RFW, and *CS* transcript, where it was positive by 5.9 to 35.6%. Overall, it is evident that N-stress decelerated all the biomass parameters and transcript levels except RL, RFW, and *CS*. While in the case of enzyme activities, the N-stress effect varied depending on the enzyme.

**Figure 8:**
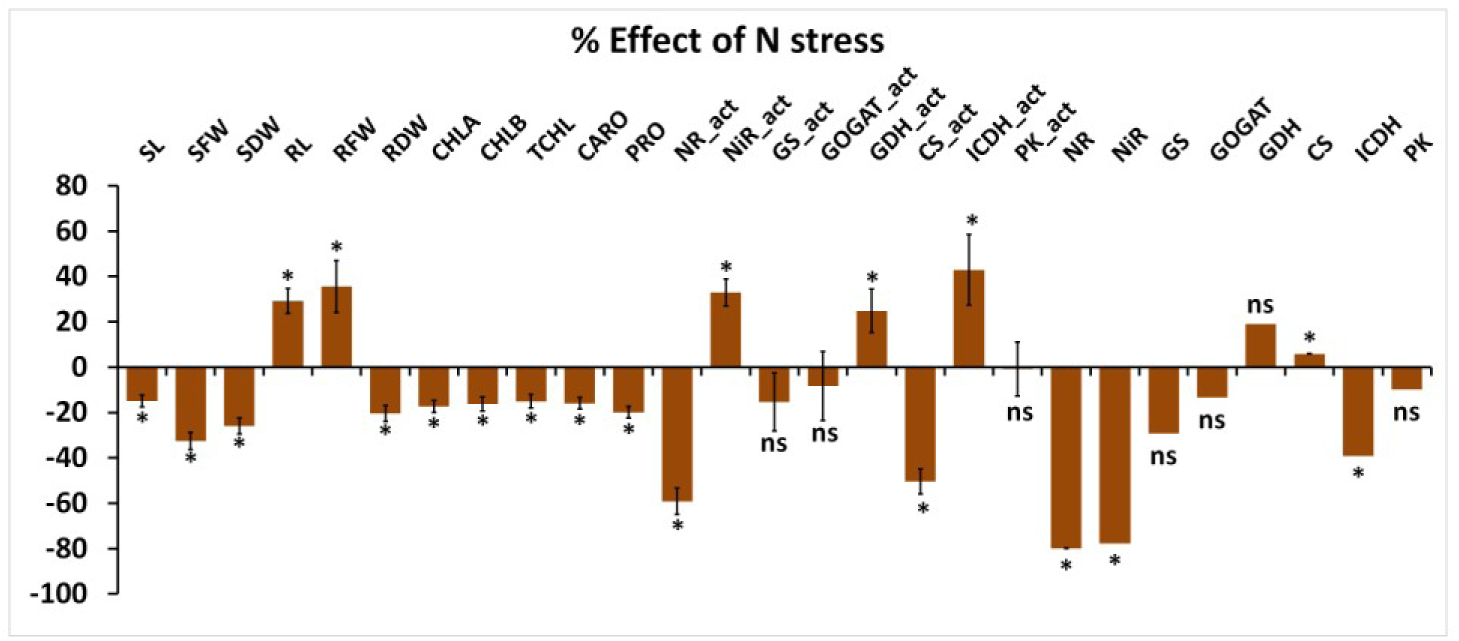
Differential effects of N on various morphological and biochemical parameters. The % effect of stressed-N (SN) minus optimum-N (ON) on each of the 19 phenotypic parameters were calculated using the mean values of 51 replicates and pooled for all the genotypes. * represents the significance at P < 0.05. PRO: protein; CHLA: Chlorophyll A; CHLB: Chlorophyll B; TCHL: Total Chlorophyll; Caro: carotenoids; SL: shoot length; SFW: shoot fresh weight, SDW: shoot dry weight, RL: root length; RFW: root fresh weight. Activities of 8 enzymes were depicted using an act suffix after the name of the enzyme, and only the enzyme name depicts its expression

### Principal Component Analysis (PCA) for morpho-physiological parameters and enzyme activities

To investigate the trait associations amongst genotypes, principal component analysis was conducted for 19 traits measured under controlled ON and SN conditions. To visualize the ratio of total variance explained by different components and variables, PCA was computed for ON and SN treatments separately (Supp File 4). Results from PCA demonstrated that under ON condition, the first five PCs explained Eigenvalues more than unity and explained a cumulative variance of 80.1%. Among these, the first two PCs such as PC-1 and PC-2, explained 26.21% and 24.54% of phenotypic variation, respectively. The major contributors (more than 30%) for PC-1 were NR, GDH, CS, and PK (Supp File 4: Supp Table 8). Likewise, PRO, SL, SFW, SDW, and RFW were highly contributed to PC-2 under ON condition (Supp File 4: Supp Table 8). Similarly, under the SN condition, the first five PCs explained Eigenvalues more than unity and explained a cumulative variance of 79.23%. Among these, the first two PCs (PC-1 and PC-2) explained 24.12% and 20.33% of the total variance, respectively. The major contributors for PC-1 were NiR, GS, GDH, and ICDH (Supp File 4: Supp Table 9). While SFW, SDW, RFW, CS, and PK (Figure 9) were found to be the major contributors to PC-2 (Figure 9). An interesting observation is that PC-1, in both cases, is driven by enzymatic activity, and PC-2 is mostly driven by morphological traits. Under both ON and SN conditions, chlorophyll and carotenoid content were found to be the major contributors to PC-1 (Figure 9).

**Figure 9:**
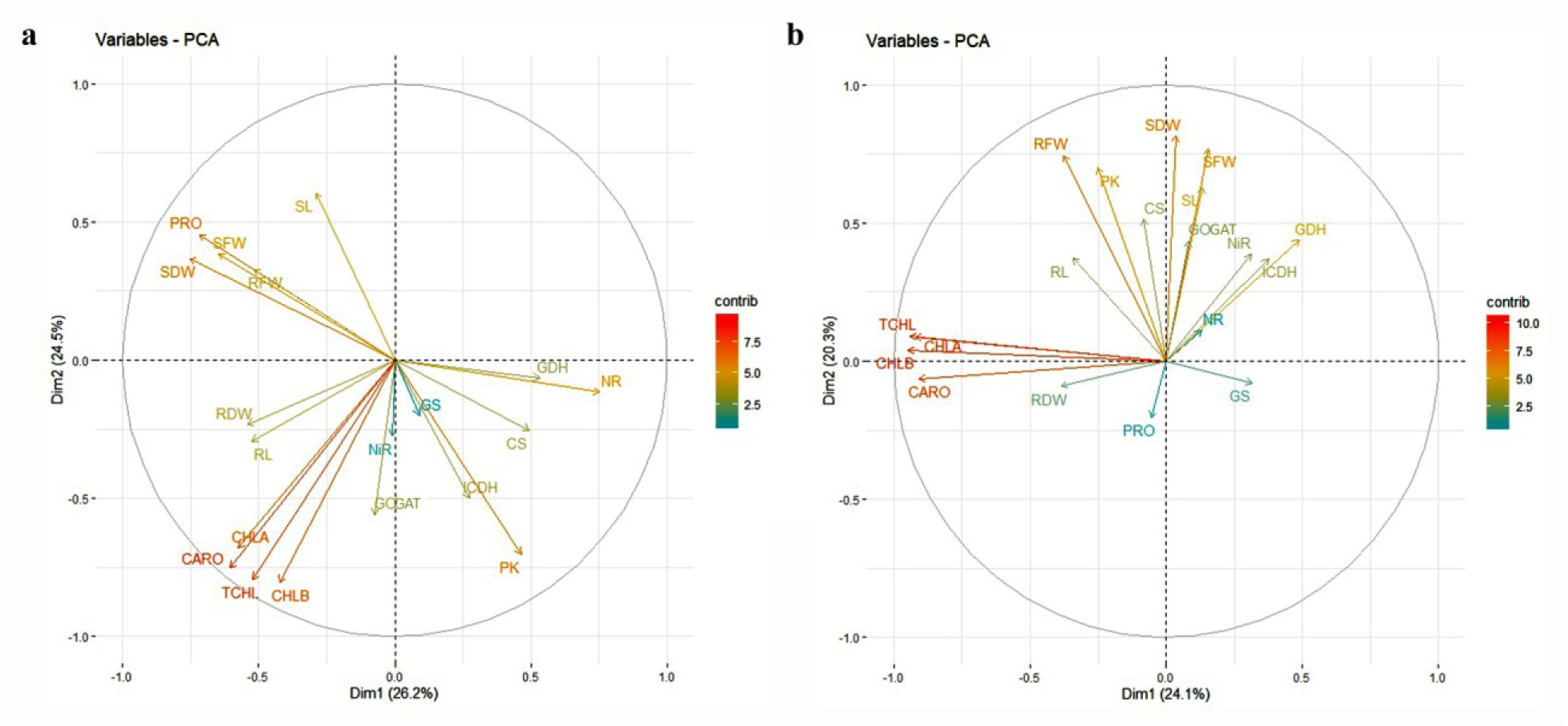
Biplot for 19 parameters measured under controlled (a) ON and (b) SN conditions.

### Correlation among different morphological parameters, enzyme activities, and expressions under ON and SN conditions

Under the ON condition, PRO was negatively correlated with *NR* and *PK* gene expressions and positively correlated with SDW (Figure 10).

**Figure 10.**
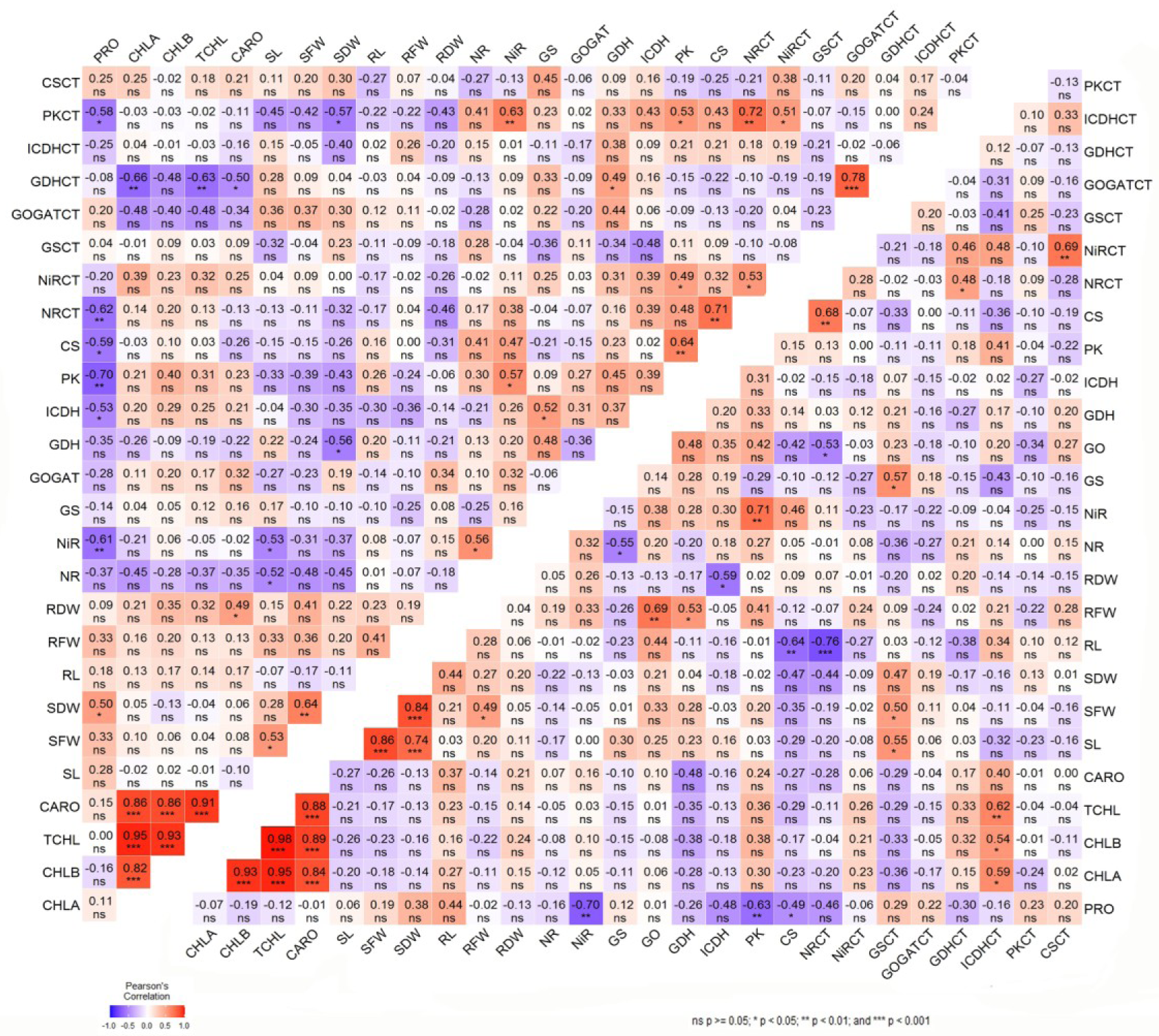
Correlation analysis among 27 parameters in hydroponics condition. The upper triangle shows correlation analysis at the ON condition, and the lower triangle shows correlation analysis at the SN condition.

Under the SN condition, a negative correlation of ICDH activity was observed with RDW. Interestingly, in the case of the N-stress condition, the root biomass and root volume increased, and hence there was a negative correlation between protein content and the root dry weight. CS activity was observed to be negatively correlated with GOGAT and RL. Shoot biomass parameters SL, SFW, and SDW were found to be positively correlated with each other. RL was observed to be positively correlated with GOGAT activity.

Out of the 8 enzymes we studied, five of them showed reduced gene expression and enzyme activity under N-stress, whereas three enzymes (NiR, CS, and ICDH) did not show a correlation between their gene expression and enzyme activity. The gene expressions of NiR and ICDH showed downregulation under N-stress, but the activities were increased.

### Analysis of Stepwise Regression

Since we are trying to find the traits affecting the output (GY), direct or indirect effects of various traits on GY were derived by stepwise backward multiple regressions under both ON and SN conditions considering the GY as the dependent variable and other traits as independent variables.

Under the ON condition, stepwise regression was performed and resulted in the sequential removal of GN, N at Head, and BM variables from the model. The remaining variables, i.e., N uptake, N at Harvest, NUtE, constitute the best-fit model and explain 94.7% of the variation in the yield (R^2^ = 0.947). Moreover, among N uptake, N at Harvest, and NUtE variables, the most contributing variable is N uptake.

Equation 1 given below shows the relationship between the GY, as the dependent variable, and other parameters as independent variables under the ON condition, and –ve and +ve signs indicate their indirect or direct effect, respectively:

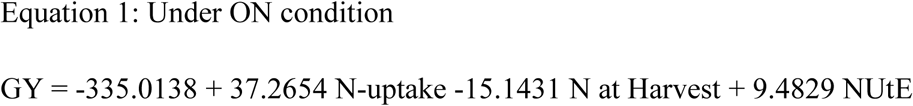

While in the case of the SN condition, N-uptake followed by BM was removed from the analysis and predicted to have no significant influence on GY. In this case, GN, N at Head, N at Harvest, and NUtE were found to have a major influence (98.4%) on GY (R^2^ = 0.984). Equation 2 below shows the relationship between the GY and other independent variables under the SN condition:

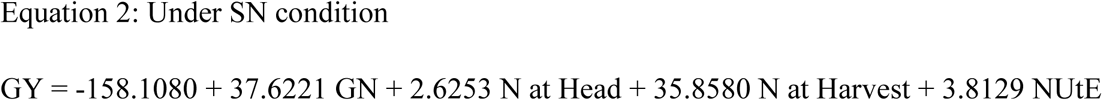

Among the hydroponics parameters, enzyme activities, protein, carotenoid, and total chlorophyll content were used as independent variables and GY as dependent variable for stepwise regression analysis. In this case, GOGAT activity, carotenoid, and total chlorophyll content were removed sequentially from the analysis showing no significant effect on GY. While rest of the enzyme activities and protein content accounted for 93.5% of the yield changes (R^2^ = 0.935) and were found to contribute maximum towards the variation influencing the GY (equation 3).

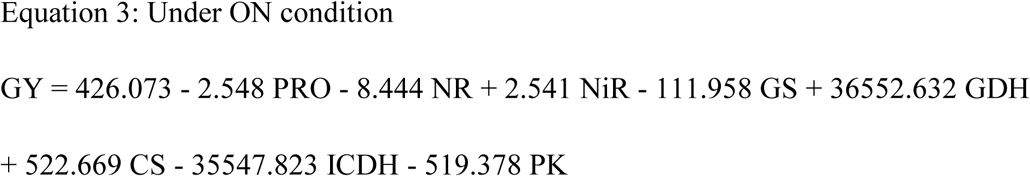

This equation shows the reason and logic very clearly. Protein content in the seedling tissue actually very important for growth and, finally, for grain yield, and the next two enzymes are NR and NiR, which are crucial for the assimilation of nitrogen and production of amino acids, and finally, proteins.

While under SN condition, the variables GOGAT activity, protein content, ICDH, CS, NR, and GS activities were removed sequentially from the stepwise regression analysis showing no significant effect on GY. The rest of the variables, i.e., Total chlorophyll, carotenoid, NiR, GDH, and PK enzyme activities (Equation 4), accounted for 60.3% of the yield changes (R^2^ = 0.603) were found to contribute maximum towards the variation influencing the GY.

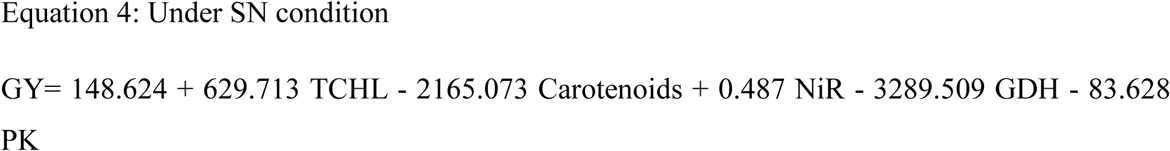

Biomass parameters measured under hydroponics conditions were also used for stepwise regression analysis, and under ON condition, SDW, RDW, RFW, and SL were removed from the analysis and predicted to have no significant effect on GY (Equation 5). While SFW and RL were found to influence the GY (R^2^ = 0.322). While under the SN condition, all the biomass parameters were removed from the analysis (Equation 6).

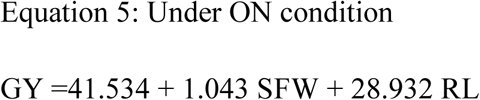

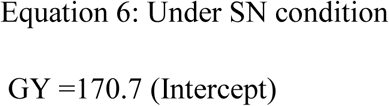

Under ON condition, the stepwise regression analysis (Equations 7 and 8) between GY and the transcripts of these eight genes showed that transcripts of *GS*, *CS*, and *GDH* were removed sequentially from the analysis, accounting for no significant effect on GY. While transcripts of *NR*, *NiR*, *GOGAT, ICDH*, and *PK* accounted for maximum contribution (Equation 7) towards the variation influencing the GY (R^2^ = 0.564).

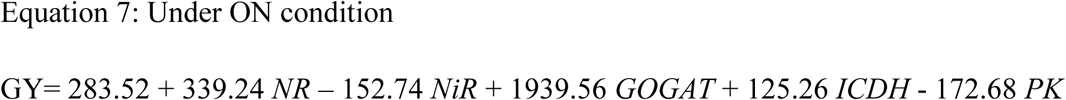

In the case of the SN condition, transcripts of *PK*, *GS*, and *ICDH* were removed from the analysis and showed no significant effect on GY. The rest of the transcripts of *NR, NiR, GOGAT, ICDH*, and *PK* accounted for maximum contribution (Equation 8) towards the variation influencing the GY (R^2^ = 0.503).

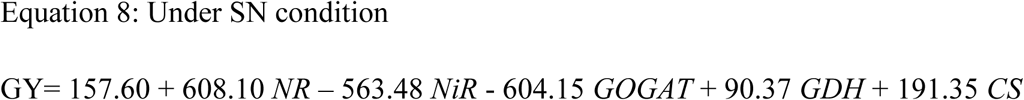

### Shortlisted parameters among field traits and hydroponics measured parameters using correlation analysis

We carried out all possible correlation analyses among all the traits and parameters which we recorded in the field and hydroponics experiment, and we tried to find the correlation between the field and hydroponics experiments. Under the ON condition, a significant positive correlation was found between N at Head and GDH (Figure 11). A negative correlation of GN and NUpE with NR, as well as NUtE with GS activity, was observed. In the case of morphological parameters, a single significant positive correlation was observed between GN and SL. We also found a significant positive correlation between GY and *GOGAT* gene expression, and a significant negative correlation between BM and *GS* gene expression was also observed. Under the SN condition, only two significant negative correlations were observed; those were between BM and RL, N at Harvest, and *GOGAT* gene expression (Figure 11).

**Figure 11.**
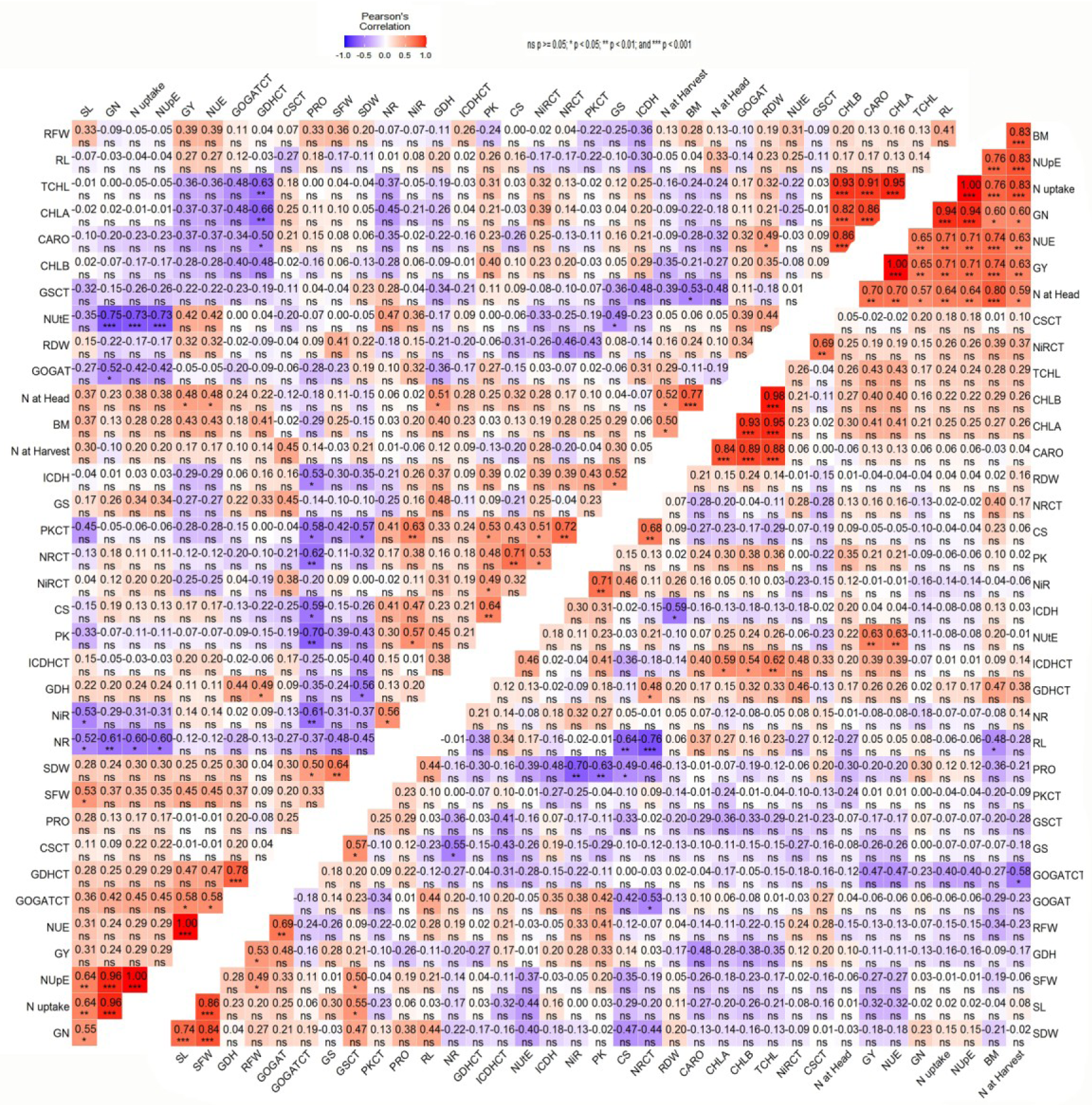
Correlation analysis among field traits and hydroponics measured parameters. The upper triangle shows correlation analysis at the ON condition, and the lower triangle shows correlation analysis at the SN condition.

## Discussion

There is ample scope for improving NUE in wheat because nitrogen loss by this important staple cereal is considerably high (Alpuerto et al., 2021; Raun & Johnson, 1999; Sylvester-Bradley & Kindred, 2009). In fact, increasing the wheat NUE is crucial to sustaining a high productivity level with a relatively low N supply (Iqbal et al., 2020; Xu et al., 2012). NUE is critically regulated by several morphological, environmental, molecular parameters and agronomic traits (Hirel et al., 2011), and a thorough understanding of these parameters/traits, along with the genotypic responses to different levels of available N in the growing media is an essential prerequisite for developing N-use efficient wheat varieties (Hajari et al., 2015). Despite various reports (Mandal et al., 2018; Raghuram et al., 2019; Sinha et al., 2015) on wheat NUE, a comprehensive study including the traits from field experiment and the various morphological, biochemical and molecular parameters under control condition has not been attempted so far. However, identification of the major traits and parameters contributing towards NUE of wheat, irrespective of the genotype, is a daunting task.

In the present study, 278 diverse wheat genotypes were analyzed for their GY and NUE-related traits under ON and NS conditions in the field, where sufficient genetic variability as well as differential responses of the different traits to the different levels of N (as shown in Figure 1 and 4) were observed in our study. Primarily, we found a positive and significant correlation of GY with all NUE-related traits viz. BM, GN, N at head, N at harvest, N uptake, NUpE, NUtE under both ON as well SN conditions. These observations highlight the importance of these traits for studying the biological NUE of wheat plant, which also reflect that these traits are mostly genotype independent. Positive and significant correlation of GY with other NUE-related traits viz. GN, HI (Harvest Index), NUpE, NUtE, and NUE were also reported by Mansour et al. 2017 and Ivić et al., 2021 and found to be consistent with our findings. However, we report that while the values of NUE-related traits alter at different levels of available soil N, these traits are correlated with each other at their respective level of soil N. NUtE was not correlated with N at Head and BM irrespective of N level, since, BM and N at Head are +vely correlated with GY and N uptake, it is not necessary that the former two traits will always be correlated with NUtE (GY/N-uptake). An inverse relation between GN and NUtE under both ON as well SN conditions may also be linked to reduced efficiency of N remobilization during the reproductive stage to the developing grain (Barraclough et al., 2010; Hawkesford, 2012; Nehe et al., 2022). For optimal use of N, it’s crucial to ensure an appropriate level of N is uptaken by the plant. Increasing N-uptake beyond the optimum level will result in diminishing returns and would not be proportionally increased in the GY and, thereby, in NUtE. The formula NUpE x NUtE = NUE highlights the inverse relationship between NUpE and NUtE. It is well established that grain protein (N) is inversely related to the yield (Geyer et al., 2022; Kramer, 1979; Laidig et al., 2017; Lawes & Gilbert, 1857; Oury et al., 2003; Oury & Godin, 2007), and that might be the probable reason why the GN is negatively correlated with NUtE. PCA from field data also suggested that among all the traits measured, N-uptake and NUpE showed their major contribution (of genetic variations available in wheat), and therefore, these traits should be given priority during the selection process targeted at wheat yield improvement. PCA also revealed that variation in N content in the plant, both after vegetative growth and at maturity (N at head and N at harvest), is more related to variation in BM production under ON as well as SN. This is an interesting indication of the important role of N acquisition in maintaining photosynthetic capacity and biomass (Gaju et al., 2016; Nehe et al., 2022; Pask et al., 2012). But for higher GY, a sufficient amount of N is required to be uptaken by the plant (Figure 12). Thus, under ON, N-uptake is higher, and N content in the plant is more positively correlated with GY. This was not the case under SN (Figure 12) because the N content in the plant was significantly low (due to low N uptake), which was mostly utilized for BM, and hence, GY was reduced significantly (Nehe et al., 2022). These observations (Figure 12) from PCA and correlation studies based on field experiment were also validated/endorsed while we carried out stepwise multiple regression.

**Figure 12:**
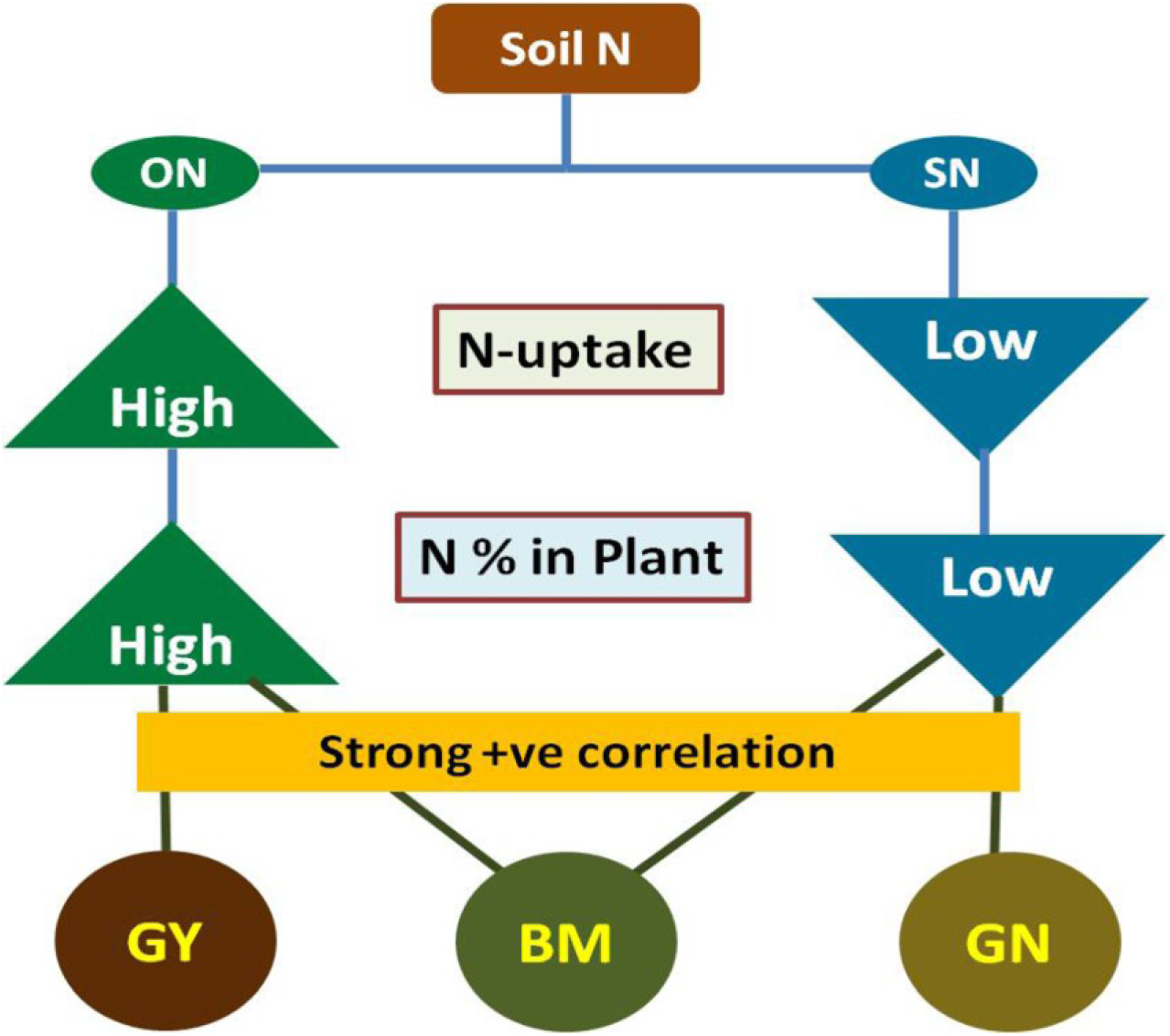
Flowchart showing the flow of N related to GY and BM. ON: Optimum-N; SN: Stressed-N; GY: Grain Yield; BM: Biomass; GN: Grain N.

Further, we tried to correlate the morphological parameters with the performance of 8 important N and C metabolizing enzymes (both enzyme activity as well as their gene expression) considering a representative set of 17 diverse genotypes, which were grown in hydroponics for precise control over N-application and environmental factors. Results showed significant variability for all the recorded parameters among the genotypes, both under ON and SN conditions (Figure 5). N-stress decelerated all the morphological parameters and photosynthetic pigment/chlorophyll and carotenoid content except RL, and RFW. While on the one hand, we found the effect of N-stress on enzymatic activity, we also observed wide genotypic variation, both under ON and SN, implying that they had diverse adaptive mechanisms to regulate N and C assimilation (Chandna et al., 2010; Gayatri et al., 2021; Sinha et al., 2015). PCA analysis clearly depicted chlorophyll contents and morphological parameters as essential factors contributing to the variations that occurred under both ON and SN conditions.

NR, GS, GOGAT, CS, and PK enzyme activity reduced significantly under SN condition suggesting their dependency on the external N concentration, while the activity of NiR, GDH, and ICDH enzymes increased under SN condition. The enhanced NiR activity (Lin et al., 2020) under N-stress condition was probably due to the reduced availability of nitrite substrate (due to lower NR activity). This increased activity of NiR, in spite of having lower gene expression of this enzyme, probably makes the plant more efficient to utilize the limited available nitrite for N assimilation, leading to enhanced nitrite reduction and ammonium assimilation. However, in spite of the presence of highly active NiR enzyme under SN condition, lack of substrate (nitrite) might produce a lower amount of product (ammonia), which affects the activity of GS and GOGAT enzymes (both in terms of reduced activities and gene expressions). The activity of the reversible enzyme GDH increased under N-stress, mainly to maintain the equilibrium of C and N metabolism. In fact, GDH works as a complementary enzyme to that of the GS/GOGAT cycle in wheat, which contributes to secondary ammonium assimilation by assimilating excess NH^+^_4_ produced due to photorespiration (Fabroni et al., 2022; Vita et al., 2018; Yan et al., 2021). Though GDH can catalyze the synthesis of glutamate under N-stress condition probably, this is catabolizing glutamate for energy production (Cammaerts & Jacobs, 1985; Miflin & Habash, 2002; Srivastava & Singh, 1987) due to the low availability of ammonia. The enhanced activity of ICDH under N-stress, in spite of lower gene expression (as we observed in case NiR), is again probably due to the low availability of substrate (citrate) produced by CS (which is a lower activity under N-stress). However, even being an active enzyme, due to the low availability of citrate under N-stress, ICDH could produce a lesser amount of 2-oxoglutarate that the plant required. There by GDH had to increase the activity to produce more 2-oxoglutarate (for TCA cycle) by catabolizing glutamate. However, the lack of correlation between the enzymatic activity and their gene expression pattern in the case of NiR, CS, and ICDH is possibly due to post-transcriptional and post-translational modifications (Gayatri et al., 2021; Kant et al., 2011).

Over all, analysis of field data with the hydroponics data revealed some interesting facts. The comprehensive analysis of field data in conjunction with hydroponics data unveiled intriguing insights and compelling patterns. NR is the rate-limiting enzyme of nitrogen metabolism and determines the plant N nutrition, and we found a significant negative correlation with NUpE and GN under ON condition as well its indirect effect on GY. Hence, NR can be used as a biochemical reference for predicting GY and GN (Croy & Hageman, 1970; F. ZHANG et al., 2015) (Li-guo et al., 2010) when N is sufficient. Since at a higher dose of N application, availability of N is more, NR activity is more and probability more of N assimilation by the plant, which might result in increased vegetative growth first and then GN followed by GY (Eilrich & Hageman, 1973). GS activity and gene expression showed an indirect effect on GY by influencing the traits like NUtE, BM (under ON), and SL (under SN). These findings confirmed that GS plays an important role in sustaining plant growth, development, and yield, as reported by Kirby et al. 2006; ZHANG et al. 2015. Moreover, the correlation of GOGAT activity with GN content under ON condition (along with NR and GS) has a straight relation with N assimilation and protein synthesis, which happened under a sufficient amount of available soil N, and this is also required for the overall physiological status of the plant (Kichey et al., 2006).

Under N-stress, correlation of root parameters (RDW and RL) and chlorophyll content was found with C-metabolizing enzymes ICDH and CS than N-assimilatory enzymes, except GDH and GOGAT. This is likely because N assimilation requires energy and resources, which may be limited under N-stress conditions. Therefore, the plant probably prioritizes the use of C-metabolism pathways to produce energy for the development of roots, which would help in N-acquisition from soil (Huang et al., 2012).

In summary, while analyzing the correlation between seedling parameters and their field traits, we observed direct and positive associations between field traits such as BM, GN, N at Head, N at Harvest, NUpE, NUtE with GY under SN conditions (Figure 13). At the seedling stage, gene expression of the *GOGAT* enzyme was found to be the most prominent parameter, which is correlated with GY directly under the ON condition and indirectly under the SN condition. Among the rest of the parameters measured at the seedling stage under hydroponics condition, SL and activities of NR, GS, GOGAT, GDH enzymes, and *GS* gene expression showed indirect relation with GY under the ON condition, while a single RL parameter has an indirect effect on GY under SN condition (Figure 13). Hence root parameters, including length, are reported to be extremely important under N-stress for foraging more nitrogen (Sinha et al., 2015).

**Figure 13:**
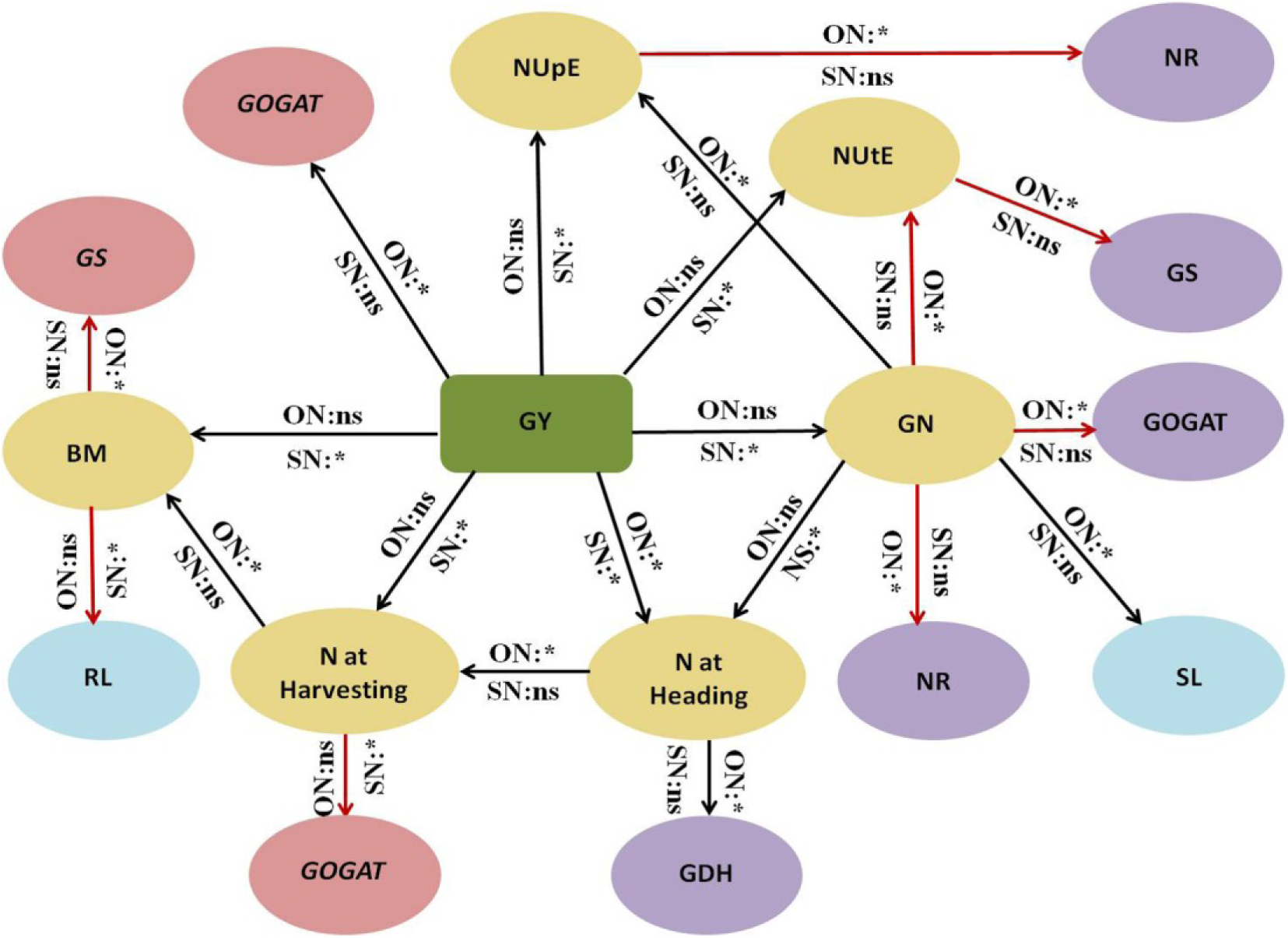
Network of seedling parameters and their NUE-related field traits. Selected hydroponics parameters were used here, which showed a significant correlation with NUE-related traits in either of the N condition (ON/SN). * denotes significant correlation, while ns denotes non-significant correlation. The black and red color lines depict the positive and negative correlations, respectively. GY: Grain Yield, BM: Biomass, GN: Grain Nitrogen, NUpE: N-uptake Efficiency, NUtE: N Utilization Efficiency, RL: Root Length, SL: Shoot Length, Enzymes names in Italics represent the expression for that particular gene, rest represent the enzyme activity

## Conclusion

Our field experiment with 278 genotypes at two levels of nitrogen revealed that BM depends on the soil nitrogen status. While under low soil nitrogen though N-uptake and GY decreases, the grain protein (GN) content increases (Figure 14). This relation is quite the opposite under a higher soil nitrogen regime, where N-uptake and GY increase but grain protein decreases. From the hydroponics experiment, we could conclude that the N-metabolizing enzymes (NR, GS, GOGAT, and GDH) have an influence on GY (Figure 14), which is not direct but mainly through N-uptake and its derived parameters. Among the seedling morphological parameters, SL is positively related to GN when nitrogen is optimum, whereas RL is negatively related to BM when nitrogen is reduced. The results of this study decipher a network of relations among the important parameters/ traits, keeping GY at the center of this network. Consequently, these traits/parameters can be considered important selection criteria for grain yield of bread wheat under reduced nitrogen application.

**Figure 14:**
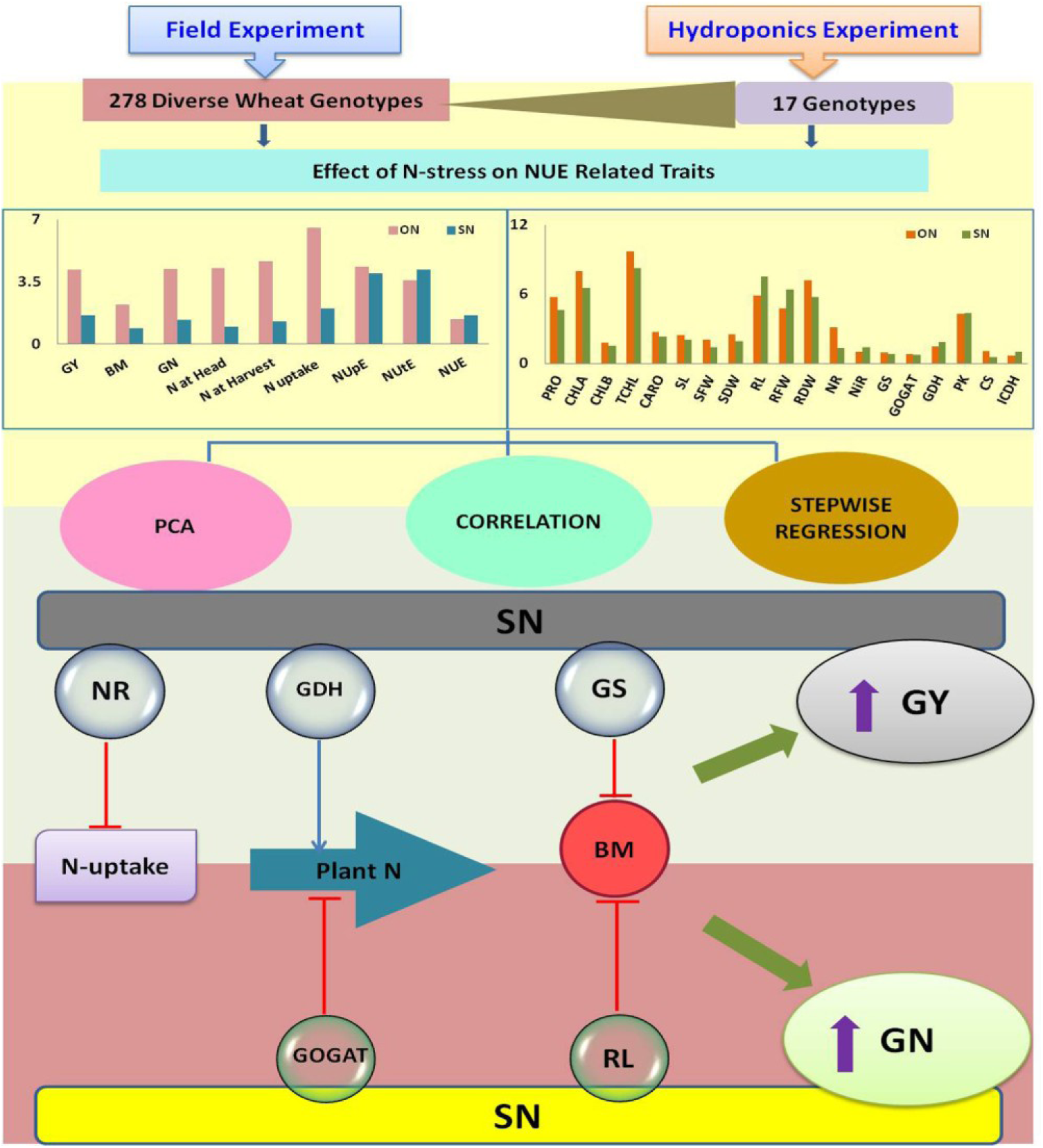
Summary of the interplay of different traits and parameters influencing GY. A set of 278 diverse wheat genotypes were evaluated in the field, and from this set, representative genotypes were further evaluated in hydroponics conditions for NUE-related traits under ON and SN conditions. All the traits and parameters were further analyzed through PCA, correlation, and stepwise regression analysis. After analysis, important traits and parameters were depicted here influencing GY. The study revealed that N-uptake and related parameters, four N-metabolizing enzymes, Nitrate Reductase (NR), Glutamine Synthetase (GS), Glutamate Synthase (GOGAT), Glutamate Dehydrogenase (GDH), and shoot as well as root length (SL, RL) parameters, had the most significant influence on grain yield. Red lines with blocks depict the –ve influence, and the blue arrow depicts the +ve influence.

## Supporting information

Supplementary.zip

## Acknowledgments

Authors would also like to acknowledge the support and guidance provided by the Project Director, ICAR-National Institute for Plant Biotechnology, New Delhi

## Authors Contribution

Conceptualization, PKM and G; Data curation, PKM; Formal analysis, G and PM; Funding acquisition, PKM; Investigation, PKM; Methodology, G; Project administration, PKM; Resources, PKM, KV; Software, G and PM; Supervision, PKM; Validation, PKM; Visualization, PKM; Writing-original draft, G; Writing review and editing PKM, KV and PM. All authors have read and agreed to the published version of the manuscript.

## References

Alpuerto, J. B., Brasier, K. G., Griffey, C. A., Thomason, W. E., & Fukao, T. (2021). Accelerated senescence and nitrogen remobilization in flag leaves enhance nitrogen use efficiency in soft red winter wheat. Plant Production Science, 24(4), 490–504.

Bargaz, A., Lyamlouli, K., Chtouki, M., Zeroual, Y., & Dhiba, D. (2018). Soil microbial resources for improving fertilizers efficiency in an integrated plant nutrient management system. Frontiers in Microbiology, 9, 1606.

Barraclough, P. B., Howarth, J. R., Jones, J., Lopez-Bellido, R., Parmar, S., Shepherd, C. E., & Hawkesford, M. J. (2010). Nitrogen efficiency of wheat: genotypic and environmental variation and prospects for improvement. European Journal of Agronomy, 33(1), 1–11.

Bharati, A., Tehlan, G., Nagar, C. K., Sinha, S. K., Venkatesh, K., & Mandal, P. K. (2022). Nitrogen stress-induced TaDof1 expression in diverse wheat genotypes and its relation with nitrogen use efficiency. Cereal Research Communications, 1–9.

Cammaerts, D., & Jacobs, M. (1985). A study of the role of glutamate dehydrogenase in the nitrogen metabolism of Arabidopsis thaliana. Planta, 163, 517–526.

Chandna, R., Gupta, S., Ahmad, A., Iqbal, M., & Prasad, M. (2010). Variability in Indian bread wheat (Triticum aestivum L.) varieties differing in nitrogen efficiency as assessed by microsatellite markers. Protoplasma, 242, 55–67.

Chardon, F., Barthélémy, J., Daniel-Vedele, F., & Masclaux-Daubresse, C. (2010). Natural variation of nitrate uptake and nitrogen use efficiency in Arabidopsis thaliana cultivated with limiting and ample nitrogen supply. Journal of Experimental Botany, 61(9), 2293–2302.

Congreves, K. A., Otchere, O., Ferland, D., Farzadfar, S., Williams, S., & Arcand, M. M. (2021). Nitrogen use efficiency definitions of today and tomorrow. Frontiers in Plant Science, 12, 637108.

Crawford, N. M., & Arst Jr, H. N. (1993). The molecular genetics of nitrate assimilation in fungi and plants. Annual Review of Genetics, 27(1), 115–146.

Croy, L. I., & Hageman, R. H. (1970). Relationship of Nitrate Reductase Activity to Grain Protein Production in Wheat 1. Crop Science, 10(3), 280–285.

Cui, Z., Zhang, H., Chen, X., Zhang, C., Ma, W., Huang, C., Zhang, W., Mi, G., Miao, Y., & Li, X. (2018). Pursuing sustainable productivity with millions of smallholder farmers. Nature, 555(7696), 363–366.

Dobermann, A. (2007). Nutrient use efficiency–measurement and management. Proceedings of the International Fertilizer Industry Association (IFA) Workshop on Fertilizer Best Management Practices, 1–28.

Dubois, F., Tercé-Laforgue, T., Gonzalez-Moro, M.-B., Estavillo, J.-M., Sangwan, R., Gallais, A., & Hirel, B. (2003). Glutamate dehydrogenase in plants: is there a new story for an old enzyme? Plant Physiology and Biochemistry, 41(6–7), 565–576.

Eilrich, G. L. t, & Hageman, R. H. (1973). Nitrate reductase activity and its relationship to accumulation of vegetative and grain nitrogen in wheat (Triticum aestivum l.) 1. Crop Science, 13(1), 59–66.

Ernst, O. R., Kemanian, A. R., Mazzilli, S., Siri-Prieto, G., & Dogliotti, S. (2020). The dos and don’ts of no-till continuous cropping: evidence from wheat yield and nitrogen use efficiency. Field Crops Research, 257, 107934.

Fabroni, S., Amenta, M., Rapisarda, S., Torrisi, B., & Licciardello, C. (2022). Amino acid metabolism and expression of genes involved in nitrogen assimilation in common oranges cv. Valencia Late. Biologia Plantarum, 66, 155–162.

Fageria, N. K., Baligar, V. C., & Li, Y. C. (2008). The role of nutrient efficient plants in improving crop yields in the twenty first century. Journal of Plant Nutrition, 31(6), 1121– 1157.

Foulkes, M. J., Hawkesford, M. J., Barraclough, P. B., Holdsworth, M. J., Kerr, S., Kightley, S., & Shewry, P. R. (2009). Identifying traits to improve the nitrogen economy of wheat: Recent advances and future prospects. Field Crops Research, 114(3), 329–342.

Foyer, C. H., Noctor, G., & Hodges, M. (2011). Respiration and nitrogen assimilation: targeting mitochondria-associated metabolism as a means to enhance nitrogen use efficiency. Journal of Experimental Botany, 62(4), 1467–1482.

Gaju, O., DeSilva, J., Carvalho, P., Hawkesford, M. J., Griffiths, S., Greenland, A., & Foulkes, M. J. (2016). Leaf photosynthesis and associations with grain yield, biomass and nitrogen-use efficiency in landraces, synthetic-derived lines and cultivars in wheat. Field Crops Research, 193, 1–15.

Gayatri, Jayaraman K., Sinha, S. K., Roy, P., & Mandal, P. K. (2021). Comparative analysis of GS2 and Fd-GOGAT genes in cultivated wheat and their progenitors under N stress. Plant Molecular Biology Reporter, 1–26.

Gayatri, Rani M., Mahato, A. K., Sinha, S. K., Dalal, M., Singh, N. K., & Mandal, P. K. (2019). Homeologue specific gene expression analysis of two vital carbon metabolizing enzymes–citrate synthase and NADP-isocitrate dehydrogenase–from wheat (Triticum aestivum L.) under nitrogen stress: homeologue specific gene expression of CS and NADP-ICDH. Applied Biochemistry and Biotechnology, 188, 569–584.

Geyer, M., Mohler, V., & Hartl, L. (2022). Genetics of the Inverse Relationship between Grain Yield and Grain Protein Content in Common Wheat. Plants, 11(16), 2146.

Giambalvo, D., Ruisi, P., Di Miceli, G., Frenda, A. S., & Amato, G. (2010). Nitrogen use efficiency and nitrogen fertilizer recovery of durum wheat genotypes as affected by interspecific competition. Agronomy Journal, 102(2), 707–715.

Good, A. G., Shrawat, A. K., & Muench, D. G. (2004). Can less yield more? Is reducing nutrient input into the environment compatible with maintaining crop production? Trends in Plant Science, 9(12), 597–605.

Guo, J., Hu, X., Gao, L., Xie, K., Ling, N., Shen, Q., Hu, S., & Guo, S. (2017). The rice production practices of high yield and high nitrogen use efficiency in Jiangsu, China. Scientific Reports, 7(1), 1–10.

Hajari, E., Snyman, S. J., & Watt, M. P. (2015). Nitrogen use efficiency of sugarcane (Saccharum spp.) varieties under in vitro conditions with varied N supply. Plant Cell, Tissue and Organ Culture (PCTOC), 122, 21–29.

Hawkesford, M. J. (2012). The diversity of nitrogen use efficiency for wheat varieties and the potential for crop improvement. Better Crops, 96(3), 10–12.

Hawkesford, M. J. (2017). Genetic variation in traits for nitrogen use efficiency in wheat. Journal of Experimental Botany, 68(10), 2627–2632.

Hawkesford, M. J., & Griffiths, S. (2019). Exploiting genetic variation in nitrogen use efficiency for cereal crop improvement. Current Opinion in Plant Biology, 49, 35–42.

Hirel, B., Tétu, T., Lea, P. J., & Dubois, F. (2011). Improving nitrogen use efficiency in crops for sustainable agriculture. Sustainability, 3(9), 1452–1485.

Huang, B., Rachmilevitch, S., & Xu, J. (2012). Root carbon and protein metabolism associated with heat tolerance. Journal of Experimental Botany, 63(9), 3455–3465.

Iqbal, A., Qiang, D., Zhun, W., Xiangru, W., Huiping, G., Hengheng, Z., Nianchang, P., Xiling, Z., & Meizhen, S. (2020). Growth and nitrogen metabolism are associated with nitrogen-use efficiency in cotton genotypes. Plant Physiology and Biochemistry, 149, 61–74.

Jain, V., Khetarpal, S., Das, R., & Abrol, Y. P. (2011). Nitrate assimilation in contrasting wheat genotypes. Physiology and Molecular Biology of Plants, 17, 137–144.

Kant, S., Bi, Y.-M., & Rothstein, S. J. (2011). Understanding plant response to nitrogen limitation for the improvement of crop nitrogen use efficiency. Journal of Experimental Botany, 62(4), 1499–1509.

Kaur, G., Asthir, B., Bains, N. S., & Farooq, M. (2015). Nitrogen nutrition, its assimilation and remobilization in diverse wheat genotypes. International Journal of Agriculture and Biology, 17(3).

Kramer, T. H. (1979). Environmental and genetic variation for protein content in winter wheat (Triticum aestivum L.). Euphytica, 28, 209–218.

Kumar, R., Taware, R., Gaur, V. S., Guru, S. K., & Kumar, A. (2009). Influence of nitrogen on the expression of TaDof1 transcription factor in wheat and its relationship with photo synthetic and ammonium assimilating efficiency. Molecular Biology Reports, 36, 2209– 2220.

Ladha, J. K., Pathak, H., Krupnik, T. J., Six, J., & van Kessel, C. (2005). Efficiency of fertilizer nitrogen in cereal production: retrospects and prospects. Advances in Agronomy, 87, 85– 156.

Laidig, F., Piepho, H.-P., Rentel, D., Drobek, T., Meyer, U., & Huesken, A. (2017). Breeding progress, environmental variation and correlation of winter wheat yield and quality traits in German official variety trials and on-farm during 1983–2014. Theoretical and Applied Genetics, 130, 223–245.

Lawes, J. B., & Gilbert, J. H. (1857). Nitrogen, on the annual yield of, per acre in different crops. RSA Journal, 6, 688.

Lea, P. J. (1993). Plant Biochemistry and Molecular Biology. In Nitrogen metabolism. Wiley, New York, NY, USA,.

Lea, P. J., Robinson, S. A., & Stewart, G. R. (1990). The biochemistry of plants: Intermediary nitrogen Metabolism. In The Biochemistry of Plants. Intermediary Nitrogen Metabolism (pp. 121–159). Academic Press, Inc.

Li-guo, F., Li-xia, S., & Huai-rui, S. (2010). Effects of exogenous NO 3-on cherry root function and enzyme activities related to nitrogen metabolism under hypoxia stress. Yingyong Shengtai Xuebao, 21(12).

Li, T., Zhang, W., Yin, J., Chadwick, D., Norse, D., Lu, Y., Liu, X., Chen, X., Zhang, F., & Powlson, D. (2018). Enhanced-sefficiency fertilizers are not a panacea for resolving the nitrogen problem. Global Change Biology, 24(2), e511–e521.

Lin, R., Bai, X., Li, F., Zhang, S., Han, L., & Xiao, K. (2020). Effects of N input level on the N-associated traits and physiological processes of winter wheat cultivated under water-saving condition. Journal of Plant Nutrition, 43(16), 2466–2479.

Mandal, V. K., Sharma, N., & Raghuram, N. (2018). Molecular targets for improvement of crop nitrogen use efficiency: Current and emerging options. Engineering Nitrogen Utilization in Crop Plants, 77–93.

Masclaux-Daubresse, C., Daniel-Vedele, F., Dechorgnat, J., Chardon, F., Gaufichon, L., & Suzuki, A. (2010). Nitrogen uptake, assimilation and remobilization in plants: challenges for sustainable and productive agriculture. Annals of Botany, 105(7), 1141–1157.

Miflin, B. J., & Habash, D. Z. (2002). The role of glutamine synthetase and glutamate dehydrogenase in nitrogen assimilation and possibilities for improvement in the nitrogen utilization of crops. Journal of Experimental Botany, 53(370), 979–987.

Moll, R. H., Kamprath, E. J., & Jackson, W. A. (1982). Analysis and interpretation of factors which contribute to efficiency of nitrogen utilization 1. Agronomy Journal, 74(3), 562–564.

Mukami, A., Ngetich, A., Mweu, C., Oduor, R. O., Muthangya, M., & Mbinda, W. M. (2019). Differential characterization of physiological and biochemical responses during drought stress in finger millet varieties. Physiology and Molecular Biology of Plants, 25, 837–846.

Nehe, A., King, J., King, I. P., Murchie, E. H., & Foulkes, M. J. (2022). Identifying variation for N-use efficiency and associated traits in amphidiploids derived from hybrids of bread wheat and the genera Aegilops, Secale, Thinopyrum and Triticum. Plos One, 17(4), e0266924.

Olivoto, T., & Lucio, A. D. (2020). metan: An R package for multi environment trial analysis. Methods in Ecology and Evolution, 11(6), 783–789.

Oury, F.-X., Berard, P., Brancourt-Hulmel, M., Heumez, E., Pluchard, P., Rousset, M., Doussinault, G., Rolland, B., Trottet, M., & Giraud, A. (2003). Yield and grain protein concentration in bread wheat: a review and a study of multi-annual data from a French breeding program [Triticum aestivum L.]. Journal of Genetics and Breeding (Italy).

Oury, F.-X., & Godin, C. (2007). Yield and grain protein concentration in bread wheat: how to use the negative relationship between the two characters to identify favourable genotypes? Euphytica, 157(1–2), 45–57.

Pask, A. J. D., Sylvester-Bradley, R., Jamieson, P. D., & Foulkes, M. J. (2012). Quantifying how winter wheat crops accumulate and use nitrogen reserves during growth. Field Crops Research, 126, 104–118.

Pathak, R. R., Jangam, A. P., Malik, A., Sharma, N., Jaiswal, D. K., & Raghuram, N. (2020). Transcriptomic and network analyses reveal distinct nitrate responses in light and dark in rice leaves (Oryza sativa Indica var. Panvel1). Scientific Reports, 10(1), 12228.

Qiao, J., Yang, L., Yan, T., Xue, F., & Zhao, D. (2012). Nitrogen fertilizer reduction in rice production for two consecutive years in the Taihu Lake area. Agriculture, Ecosystems & Environment, 146(1), 103–112.

Quevedo-Amaya, Y. M., Beltrán-Medina, J. I., Hoyos-Cartagena, J. Á., Calderón-Carvajal, J. E., & Barragán-Quijano, E. (2020). Selection of sowing date and biofertilization as alternatives to improve the yield and profitability of the F68 rice variety. Agronomía Colombiana, 38(1), 61–72.

Raghuram, N., & Sharma, N. (2019). Comprehensive biotechnology (Vol. 4, ed). Elsevier. https://doi.org/doi:10.1016/b978-0-444-64046-8.00222-6

Raun, W. R., & Johnson, G. V. (1999). Improving nitrogen use efficiency for cereal production. Agronomy Journal, 91(3), 357–363.

Santiago-Arenas, R., Fanshuri, B. A., Hadi, S. N., Ullah, H., & Datta, A. (2020). Nitrogen fertiliser and establishment method affect growth, yield and nitrogen use efficiency of rice under alternate wetting and drying irrigation. Annals of Applied Biology, 176(3), 314–327.

Scheible, W.-R., Gonzalez-Fontes, A., Lauerer, M., Muller-Rober, B., Caboche, M., & Stitt, M. (1997). Nitrate acts as a signal to induce organic acid metabolism and repress starch metabolism in tobacco. The Plant Cell, 9(5), 783–798.

Scheible, W. R., Krapp, A., & Stitt, M. (2000). Reciprocal diurnal changes of phosphoenolpyruvate carboxylase expression and cytosolic pyruvate kinase, citrate synthase and NADP isocitrate dehydrogenase expression regulate organic acid metabolism during nitrate assimilation in tobacco leaves. *Plant*, Cell & Environment, 23(11), 1155–1167.

Schmittgen, T. D., & Livak, K. J. (2008). Analyzing real-time PCR data by the comparative CT method. Nature Protocols, 3(6), 1101–1108.

Sevanthi, A. M., Sinha, S. K., Rani, M., Saini, M. R., Kumari, S., Kaushik, M., Prakash, C., Singh, G. P., Mohapatra, T., & Mandal, P. K. (2021). Integration of dual stress transcriptomes and major QTLs from a pair of genotypes contrasting for drought and chronic nitrogen starvation identifies key stress responsive genes in rice. Rice, 14(1), 1–28.

Shrawat, A. K., & Good, A. G. (2008). Genetic engineering approaches to improving nitrogen use efficiency. ISB News Report, 1–5.

Sieling, K., & Kage, H. (2008). The potential of semi-dwarf oilseed rape genotypes to reduce the risk of N leaching. The Journal of Agricultural Science, 146(1), 77–84.

Sinha, S. K., Rani, M., Bansal, N., Gayatri, Venkatesh K., & Mandal, P. K. (2015). Nitrate starvation induced changes in root system architecture, carbon: nitrogen metabolism, and miRNA expression in nitrogen-responsive wheat genotypes. Applied Biochemistry and Biotechnology, 177, 1299–1312.

Sinha, V. B., Jangam, A. P., & Raghuram, N. (2020). Biological determinants of crop nitrogen use efficiency and biotechnological avenues for improvement. Just Enough Nitrogen: Perspectives on How to Get There for Regions with Too Much and Too Little Nitrogen, 157–171.

Srivastava, H. S., & Singh, R. P. (1987). Role and regulation of L-glutamate dehydrogenase activity in higher plants. Phytochemistry, 26(3), 597–610.

Stitt, M., & Krapp, A. (1999). The interaction between elevated carbon dioxide and nitrogen nutrition: the physiological and molecular background. *Plant*, Cell & Environment, 22(6), 583–621.

Suzuki, A., & Knaff, D. B. (2005). Glutamate synthase: structural, mechanistic and regulatory properties, and role in the amino acid metabolism. Photosynthesis Research, 83, 191–217.

Sylvester-Bradley, R., & Kindred, D. R. (2009). Analysing nitrogen responses of cereals to prioritize routes to the improvement of nitrogen use efficiency. Journal of Experimental Botany, 60(7), 1939–1951.

Tang, W., Ye, J., Yao, X., Zhao, P., Xuan, W., Tian, Y., Zhang, Y., Xu, S., An, H., & Chen, G. (2019). Genome-wide associated study identifies NAC42-activated nitrate transporter conferring high nitrogen use efficiency in rice. Nature Communications, 10(1), 5279.

Tercé-Laforgue, T., Dubois, F., Ferrario-Méry, S., Crecenzo, M. P. D., Sangwan, R., & Hirel, B. (2004). Glutamatedehydrogenase of tobacco (Nicotiana tabacum L.) is mainly induced in the cytosol of phloem companion cells whenammonia is provided either externally or released during photorespiration. Plant Physiol, 136, 4308–4317.

Vita, F., Giuntoli, B., Arena, S., Quaranta, F., Bertolini, E., Lucarotti, V., Guglielminetti, L., Alessio, M., Scaloni, A., & Alpi, A. (2018). Effects of different nitrogen fertilizers on two wheat cultivars: An integrated approach. Plant Direct, 2(10), e00089.

Wickham, H. (2009). ggplot2: elegant graphics for data analysis New York. NY: Springer.

Xu, G., Fan, X., & Miller, A. J. (2012). Plant nitrogen assimilation and use efficiency. Annual Review of Plant Biology, 63, 153–182.

Yan, L. U., Gong, Y., Luo, Q., Dai, G.-X., Teng, Z., He, Y., Wu, X., Liu, C., Tang, D., & Ye, N. (2021). Heterologous expression of fungal AcGDH alleviates ammonium toxicity and suppresses photorespiration, thereby improving drought tolerance in rice. Plant Science, 305, 110769.

Yanagisawa, S. (2002). The Dof family of plant transcription factors. Trends in Plant Science, 7(12), 555–560.

Yanagisawa, S. (2004). Dof domain proteins: plant-specific transcription factors associated with diverse phenomena unique to plants. Plant and Cell Physiology, 45(4), 386–391.

Zhang, F., Si, G. A. O., Zhao, Y., Zhao, X., Liu, X., & Kai, X. (2015). Growth traits and nitrogen assimilation-associated physiological parameters of wheat (Triticum aestivum L.) under low and high N conditions. Journal of Integrative Agriculture, 14(7), 1295–1308.

Zhang, H., Fu, X., Wang, X., Gui, H., Dong, Q., Pang, N., Wang, Z., Zhang, X., & Song, M. (2018). Identification and screening of nitrogen-efficient cotton genotypes under low and normal nitrogen environments at the seedling stage. Journal of Cotton Research, 1(1), 1–11.

Zheng, Z.-L. (2009). Carbon and nitrogen nutrient balance signaling in plants. Plant Signaling & Behavior, 4(7), 584–591.

