## Supplementary.zip for "Unraveling the Interplay of Different Traits and Parameters Related to Nitrogen Use Efficiency in Wheat: Insights for Grain Yield Influence": Suppl File 1.docx

| **Traits** | **PC-1** | **PC-2** | **PC-3** | **PC-4** | **PC-5** |
| --- | --- | --- | --- | --- | --- |
| **GY** | 0.32022 | 0.376 | 0.325952 | 0.308618 | -0.03482 |
| **BM** | 0.33658 | 0.270636 | -0.12451 | -0.45613 | 0.76803 |
| **GN** | 0.349748 | -0.35507 | 0.311069 | -0.03487 | -0.00086 |
| **N at Head** | 0.31368 | 0.287442 | -0.10114 | -0.63782 | -0.63398 |
| **N at Harvest** | 0.286605 | 0.126683 | -0.81301 | 0.394324 | -0.0678 |
| **N-uptake** | 0.408188 | -0.27366 | 0.003184 | 0.101808 | -0.02361 |
| **NUpE** | 0.408188 | -0.27366 | 0.003184 | 0.101808 | -0.02361 |
| **NUtE** | -0.21339 | 0.51911 | 0.063266 | 0.131161 | -0.00785 |
| **NUE** | 0.32022 | 0.376 | 0.325952 | 0.308618 | -0.03482 |
| **eigen value** | 4.731576 | 2.784703 | 1.610694 | 1.211916 | 0.578551 |
| **variance** | 43.01432 | 25.31548 | 14.64267 | 11.01741 | 5.259552 |
| **cumulative.variance.percent** | 43.01432 | 68.3298 | 82.97247 | 93.98989 | 99.24944 |

**Supp Table 5: Principal component analysis of 9 NUE traits measured at field under ON conditions.** Keys: GY: grain yield; BM: biomass, GN: grain nitrogen, NUpE: Nitrogen Uptake efficiency; NUtE: Nitrogen Utilization Efficiency; NUE: Nitrogen Use Efficiency

| **Traits** | **PC-1** | **PC-2** | **PC-3** | **PC-4** | **PC-5** |
| --- | --- | --- | --- | --- | --- |
| **GY** | 0.345096 | -0.39499 | 0.0647 | -0.06434 | 0.053056 |
| **BM** | 0.352397 | 0.098882 | -0.40462 | 0.012124 | -0.83761 |
| **GN** | 0.351215 | 0.06641 | 0.532488 | 0.229202 | -0.09146 |
| **N at Head** | 0.323429 | 0.114709 | -0.47598 | 0.709004 | 0.390466 |
| **N at Harvest** | 0.32987 | 0.206918 | -0.4027 | -0.65307 | 0.352443 |
| **N-uptake** | 0.387732 | 0.127096 | 0.251932 | -0.07006 | 0.062126 |
| **NUpE** | 0.387732 | 0.127096 | 0.251932 | -0.07006 | 0.062126 |
| **NUtE** | -0.01377 | -0.76517 | -0.16943 | -0.00571 | 0.003524 |
| **NUE** | 0.345096 | -0.39499 | 0.0647 | -0.06434 | 0.053056 |
| **eigen value** | 6.11669 | 1.629535 | 0.772185 | 0.283368 | 0.139654 |
| **variance** | 67.96322 | 18.10594 | 8.579835 | 3.148536 | 1.551716 |
| **cumulative.variance.percent** | 67.96322 | 86.06917 | 94.649 | 97.79754 | 99.34925 |

**Supp Table 6: Principal component analysis of 9 NUE traits measured at field under SN conditions.** Keys: GY: grain yield; BM: biomass, GN: grain nitrogen, NUpE: Nitrogen Uptake efficiency; NUtE: Nitrogen Utilization Efficiency; NUE: Nitrogen Use Efficiency
