## Supplementary.zip for "Unraveling the Interplay of Different Traits and Parameters Related to Nitrogen Use Efficiency in Wheat: Insights for Grain Yield Influence": Suppl File 2.docx

BM (g/m2)

| Genotypes | BM (g/m2) |
| --- | --- |
| 1 | 1150 |
| 2 | 856.547 |
| 3 | 1454.17 |
| 4 | 1078.33 |
| 5 | 847.5 |
| 6 | 868.333 |
| 7 | 1258.33 |
| 8 | 1213.33 |
| 9 | 1337.5 |
| 10 | 1333.33 |
| 11 | 1391.67 |
| 12 | 955 |
| 13 | 1190 |
| 14 | 942.5 |
| 15 | 850 |
| 16 | 920 |
| 17 | 898.571 |
| 18 | 955 |
| 19 | 1113.04 |
| 20 | 758.333 |
| 21 | 862.778 |
| 22 | 1151.67 |
| 23 | 989.583 |
| 24 | 1091.67 |
| 25 | 1365 |
| 26 | 1025 |
| 27 | 1054.29 |
| 28 | 1157.92 |
| 29 | 1017.5 |
| 30 | 1090 |
| 31 | 1333.33 |
| 32 | 901.19 |
| 33 | 979.167 |
| 34 | 836.111 |
| 35 | 877.791 |
| 36 | 1038.57 |
| 37 | 1008.57 |
| 38 | 871.667 |
| 39 | 1337.89 |
| 40 | 1156.67 |
| 41 | 965.476 |
| 42 | 962.857 |
| 43 | 797.619 |
| 44 | 933.333 |
| 45 | 1111.67 |
| 46 | 925 |
| 47 | 897.321 |
| 48 | 870 |
| 49 | 970 |
| 50 | 1273.04 |
| 51 | 1308.33 |
| 52 | 986.667 |
| 53 | 1690 |
| 54 | 1142.35 |
| 55 | 1140 |
| 56 | 913.095 |
| 57 | 932.143 |
| 58 | 1150 |
| 59 | 963.393 |
| 60 | 1345 |
| 61 | 702.652 |
| 62 | 901.667 |
| 63 | 927.976 |
| 64 | 658.081 |
| 65 | 1360 |
| 66 | 1186.61 |
| 67 | 1807.35 |
| 68 | 1058.11 |
| 69 | 1096.97 |
| 70 | 847.715 |
| 71 | 1440.57 |
| 72 | 948.572 |
| 73 | 1338.07 |
| 74 | 656.832 |
| 75 | 1170.5 |
| 76 | 1501.28 |
| 77 | 1100 |
| 78 | 981.093 |
| 79 | 1039.39 |
| 80 | 1377.38 |
| 81 | 1091.67 |
| 82 | 1035.12 |
| 83 | 1210.92 |
| 84 | 1087.5 |
| 85 | 1023.33 |
| 86 | 1000 |
| 87 | 1058.33 |
| 88 | 880.159 |
| 89 | 854.167 |
| 90 | 921.667 |
| 91 | 954.464 |
| 92 | 1264.39 |
| 93 | 1224.17 |
| 94 | 1691.79 |
| 95 | 758.273 |
| 96 | 1601.67 |
| 97 | 876.25 |
| 98 | 783.036 |
| 99 | 963.036 |
| 100 | 836.397 |
| 101 | 988.661 |
| 102 | 862.5 |
| 103 | 1248.93 |
| 104 | 1153.07 |
| 105 | 1028.33 |
| 106 | 1095.24 |
| 107 | 1038.33 |
| 108 | 1093.9 |
| 109 | 1311.4 |
| 110 | 953.066 |
| 111 | 991.667 |
| 112 | 1048.57 |
| 113 | 1234.29 |
| 114 | 1273.15 |
| 115 | 952.679 |
| 116 | 1111.4 |
| 117 | 815.893 |
| 118 | 1119.76 |
| 119 | 785.536 |
| 120 | 1266.67 |
| 121 | 1527.5 |
| 122 | 1163.21 |
| 123 | 1542.5 |
| 124 | 1426.67 |
| 125 | 1015.57 |
| 126 | 1306.79 |
| 127 | 1433.75 |
| 128 | 1331.43 |
| 129 | 1172.41 |
| 130 | 1136.4 |
| 131 | 732.143 |
| 132 | 923.036 |
| 133 | 952.47 |
| 134 | 1330.54 |
| 135 | 1211.4 |
| 136 | 1427.08 |
| 137 | 856.607 |
| 138 | 879.286 |
| 139 | 1410 |
| 140 | 1405 |
| 141 | 1275 |
| 142 | 1155 |
| 143 | 998.898 |
| 144 | 985.714 |
| 145 | 1398.9 |
| 146 | 1163.07 |
| 147 | 1008.33 |
| 148 | 908.898 |
| 149 | 1067.35 |
| 150 | 1192.44 |
| 151 | 978.065 |
| 152 | 991.667 |
| 153 | 1100.57 |
| 154 | 848.899 |
| 155 | 998.899 |
| 156 | 905.578 |
| 157 | 1423.04 |
| 158 | 1310 |
| 159 | 1120 |
| 160 | 1286.38 |
| 161 | 1046.25 |
| 162 | 1090 |
| 163 | 1050.54 |
| 164 | 1223.88 |
| 165 | 1130.13 |
| 166 | 1418.04 |
| 167 | 1115.13 |
| 168 | 1485.36 |
| 169 | 1285 |
| 170 | 1344.91 |
| 171 | 868.81 |
| 172 | 1116.67 |
| 173 | 1373.89 |
| 174 | 1465.58 |
| 175 | 840.178 |
| 176 | 1286.61 |
| 177 | 1078.33 |
| 178 | 1545.58 |
| 179 | 1245.83 |
| 180 | 1026.67 |
| 181 | 1376.67 |
| 182 | 1342.64 |
| 183 | 1104.17 |
| 184 | 1199.24 |
| 185 | 1285.58 |
| 186 | 1328.08 |
| 187 | 1190.57 |
| 188 | 1048.89 |
| 189 | 1358.43 |
| 190 | 1545 |
| 191 | 1586.39 |
| 192 | 1153.07 |
| 193 | 1261.39 |
| 194 | 1576.67 |
| 195 | 1825 |
| 196 | 1223.89 |
| 197 | 1655.57 |
| 198 | 1265.54 |
| 199 | 1350 |
| 200 | 901.667 |

**Supp Figure 1:** Biomass (BM) of representative 17 genotypes selected from a diverse set of 278 wheat genotypes. Red lines indicate the selected 17 genotypes used for the hydroponics experiment

GY (g/m2)

|  | GY/P (g/m2) |
| --- | --- |
| 1 | 406.4 |
| 2 | 358.732 |
| 3 | 530.458 |
| 4 | 450.083 |
| 5 | 330.9 |
| 6 | 414.9 |
| 7 | 404.333 |
| 8 | 369.65 |
| 9 | 420.583 |
| 10 | 405.167 |
| 11 | 401.03 |
| 12 | 417.892 |
| 13 | 458.6 |
| 14 | 425.565 |
| 15 | 349.428 |
| 16 | 413.75 |
| 17 | 368.36 |
| 18 | 408.875 |
| 19 | 394.744 |
| 20 | 312.429 |
| 21 | 344.778 |
| 22 | 492.183 |
| 23 | 429.458 |
| 24 | 434.883 |
| 25 | 545.85 |
| 26 | 407.125 |
| 27 | 356.214 |
| 28 | 399.737 |
| 29 | 405.542 |
| 30 | 285.075 |
| 31 | 433.067 |
| 32 | 365.238 |
| 33 | 390.125 |
| 34 | 367.513 |
| 35 | 401.807 |
| 36 | 404.7 |
| 37 | 382.871 |
| 38 | 380.815 |
| 39 | 455.031 |
| 40 | 452.917 |
| 41 | 331.547 |
| 42 | 326.871 |
| 43 | 292.547 |
| 44 | 371.155 |
| 45 | 406.75 |
| 46 | 359.1 |
| 47 | 359.092 |
| 48 | 306.7 |
| 49 | 429.4 |
| 50 | 481.945 |
| 51 | 549.172 |
| 52 | 416.333 |
| 53 | 588.075 |
| 54 | 458.082 |
| 55 | 462.829 |
| 56 | 450.024 |
| 57 | 399.809 |
| 58 | 422.917 |
| 59 | 400.146 |
| 60 | 601.833 |
| 61 | 243.138 |
| 62 | 376.65 |
| 63 | 350.539 |
| 64 | 274.762 |
| 65 | 501.775 |
| 66 | 427.384 |
| 67 | 606.638 |
| 68 | 409.473 |
| 69 | 472.722 |
| 70 | 352.35 |
| 71 | 488.272 |
| 72 | 338.307 |
| 73 | 550.57 |
| 74 | 271.202 |
| 75 | 409.542 |
| 76 | 523.556 |
| 77 | 465.583 |
| 78 | 393.334 |
| 79 | 447.765 |
| 80 | 518.315 |
| 81 | 444.467 |
| 82 | 456.25 |
| 83 | 475.639 |
| 84 | 495.325 |
| 85 | 424.65 |
| 86 | 396.317 |
| 87 | 457.5 |
| 88 | 341.837 |
| 89 | 355.708 |
| 90 | 400.617 |
| 91 | 407.223 |
| 92 | 535.148 |
| 93 | 505.374 |
| 94 | 705.868 |
| 95 | 311.759 |
| 96 | 390.25 |
| 97 | 320.637 |
| 98 | 303.732 |
| 99 | 350.713 |
| 100 | 308.062 |
| 101 | 394.062 |
| 102 | 340.125 |
| 103 | 500.814 |
| 104 | 477.81 |
| 105 | 457.984 |
| 106 | 401.286 |
| 107 | 346.492 |
| 108 | 443.48 |
| 109 | 424.947 |
| 110 | 337.667 |
| 111 | 391.317 |
| 112 | 413.114 |
| 113 | 506.179 |
| 114 | 509.213 |
| 115 | 418.465 |
| 116 | 526.6 |
| 117 | 266.143 |
| 118 | 373.969 |
| 119 | 321.208 |
| 120 | 367.917 |
| 121 | 627.05 |
| 122 | 458.804 |
| 123 | 489.809 |
| 124 | 439.667 |
| 125 | 370.28 |
| 126 | 522.482 |
| 127 | 492.398 |
| 128 | 566.969 |
| 129 | 318.854 |
| 130 | 325.655 |
| 131 | 342.617 |
| 132 | 400.914 |
| 133 | 372.065 |
| 134 | 525.845 |
| 135 | 459.847 |
| 136 | 537.542 |
| 137 | 377.904 |
| 138 | 365.489 |
| 139 | 398.8 |
| 140 | 383.875 |
| 141 | 387.367 |
| 142 | 481.842 |
| 143 | 371.673 |
| 144 | 438.686 |
| 145 | 572.446 |
| 146 | 480.811 |
| 147 | 357.333 |
| 148 | 283.902 |
| 149 | 418.223 |
| 150 | 442.239 |
| 151 | 360.797 |
| 152 | 419.521 |
| 153 | 363.514 |
| 154 | 359.733 |
| 155 | 371.028 |
| 156 | 367.174 |
| 157 | 399.329 |
| 158 | 561.3 |
| 159 | 317.2 |
| 160 | 539.952 |
| 161 | 415.67 |
| 162 | 419.425 |
| 163 | 406.842 |
| 164 | 496.625 |
| 165 | 385.575 |
| 166 | 649.583 |
| 167 | 312.969 |
| 168 | 456.618 |
| 169 | 391.075 |
| 170 | 447.429 |
| 171 | 356.467 |
| 172 | 451.25 |
| 173 | 583.867 |
| 174 | 589.977 |
| 175 | 375.6 |
| 176 | 322.717 |
| 177 | 294.189 |
| 178 | 448.667 |
| 179 | 375.483 |
| 180 | 361.075 |
| 181 | 481.917 |
| 182 | 423.274 |
| 183 | 322.006 |
| 184 | 304.254 |
| 185 | 328.889 |
| 186 | 320.074 |
| 187 | 360.551 |
| 188 | 319.624 |
| 189 | 392.034 |
| 190 | 409.95 |
| 191 | 475.143 |
| 192 | 495.567 |
| 193 | 370.237 |
| 194 | 522.66 |
| 195 | 527.25 |
| 196 | 412.296 |
| 197 | 377.487 |
| 198 | 318.627 |
| 199 | 398.25 |
| 200 | 263.083 |

**Supp Figure 2:** Grain Yield (GY) of representative 17 genotypes selected from a diverse set of 278 wheat genotypes. Red lines indicate the selected 17 genotypes used for the hydroponics experiment
