## Supplementary.zip for "Unraveling the Interplay of Different Traits and Parameters Related to Nitrogen Use Efficiency in Wheat: Insights for Grain Yield Influence": Suppl File 4.docx

| **Parameters** | **PC-1** | **PC-2** | **PC-3** | **PC-4** | **PC-5** |
| --- | --- | --- | --- | --- | --- |
| **PRO** | -0.71914 | 0.454353 | 0.001767 | 0.128379 | -0.35133 |
| **CHLA** | -0.5758 | -0.67933 | 0.188917 | -0.22011 | -0.14811 |
| **CHLB** | -0.42393 | -0.80456 | 0.114523 | -0.24022 | 0.05757 |
| **TCHL** | -0.52197 | -0.79389 | 0.162062 | -0.18644 | -0.0811 |
| **CARO** | -0.60553 | -0.7519 | 0.046018 | 0.065342 | -0.02383 |
| **SL** | -0.28889 | 0.604884 | 0.434581 | -0.33379 | 0.258842 |
| **SFW** | -0.64815 | 0.385343 | -0.00188 | -0.23003 | 0.544793 |
| **SDW** | -0.75347 | 0.366702 | 0.040012 | -0.32985 | 0.104383 |
| **RL** | -0.52592 | -0.29305 | 0.006842 | -0.04989 | -0.41588 |
| **RFW** | -0.51741 | 0.32871 | -0.29856 | 0.01982 | 0.349738 |
| **RDW** | -0.54274 | -0.23336 | -0.37111 | 0.280646 | 0.503956 |
| **NR** | 0.753927 | -0.11655 | -0.46722 | -0.10349 | -0.06263 |
| **NiR** | -0.00801 | -0.27457 | -0.77057 | 0.012724 | 0.333666 |
| **GS** | 0.089841 | -0.20048 | 0.612908 | 0.446874 | 0.32894 |
| **GOGAT** | -0.07536 | -0.5587 | -0.3987 | 0.421468 | 0.084346 |
| **GDH** | 0.534306 | -0.06395 | 0.49036 | -0.08056 | 0.367286 |
| **CS** | 0.494328 | -0.25347 | -0.12212 | -0.75124 | 0.089537 |
| **ICDH** | 0.274181 | -0.5003 | 0.529812 | 0.299844 | 0.210453 |
| **PK** | 0.464634 | -0.70346 | -0.02731 | -0.32238 | 0.216262 |
| **eigen value** | 4.98 | 4.66 | 2.38 | 1.67 | 1.54 |
| **variance %** | 26.2 | 24.5 | 12.5 | 8.79 | 8.13 |
| **cumulative variance %** | 26.21481 | 50.75632 | 63.27245 | 72.05841 | 80.18639 |

**Supp Table 8: Principal component analysis of the measured morpho-physiological parameters and enzyme assays under ON conditions.** PRO: protein; CHLA: chlorophyll A; CHLB: chlorophyll B; TCHL: total chlorophyll; Caro: carotenoids; SL: shoot length; SFW: shoot fresh weight, SDW: shoot dry weight, RL: root length; RFW: root fresh weight

| **Parameters** | **PC-1** | **PC-2** | **PC-3** | **PC-4** | **PC-5** |
| --- | --- | --- | --- | --- | --- |
| **PRO** | -0.054 | -0.20512 | -0.81589 | -0.00926 | 0.401098 |
| **CHLA** | -0.92181 | 0.088 | 0.172539 | -0.24165 | 0.104984 |
| **CHLB** | -0.94815 | 0.040994 | 0.227938 | -0.1467 | -0.00877 |
| **TCHL** | -0.94068 | 0.091141 | 0.173876 | -0.1747 | 0.098563 |
| **CARO** | -0.90861 | -0.06282 | 0.028575 | -0.22243 | 0.199646 |
| **SL** | 0.133167 | 0.629549 | -0.32994 | -0.35755 | -0.07483 |
| **SFW** | 0.153233 | 0.769062 | -0.57971 | -0.05877 | -0.11833 |
| **SDW** | 0.033827 | 0.816392 | -0.4778 | 0.003228 | -0.17736 |
| **RL** | -0.34287 | 0.373144 | -0.25677 | 0.457536 | 0.222917 |
| **RFW** | -0.37615 | 0.745049 | -0.39709 | 0.07116 | -0.04452 |
| **RDW** | -0.38505 | -0.09054 | -0.06579 | 0.285289 | -0.4397 |
| **NR** | 0.130362 | 0.111642 | 0.166075 | 0.752941 | 0.457171 |
| **NiR** | 0.313682 | 0.386471 | 0.710079 | 0.023551 | -0.05692 |
| **GS** | 0.316701 | -0.0793 | -0.03115 | -0.80214 | 0.181974 |
| **GO** | 0.082389 | 0.435691 | 0.104342 | -0.10499 | 0.787076 |
| **GDH** | 0.487815 | 0.439253 | 0.254697 | -0.21522 | 0.023939 |
| **CS** | -0.08257 | 0.513053 | 0.382642 | -0.02791 | -0.44428 |
| **ICDH** | 0.375369 | 0.370396 | 0.545298 | -0.16701 | 0.280871 |
| **PK** | -0.25127 | 0.700084 | 0.531829 | 0.223909 | 0.037003 |
| **eigen value** | 4.58 | 3.86 | 3.01 | 1.93 | 1.66 |
| **variance %** | 24.12 | 20.33 | 15.87 | 10.18 | 8.74 |
| **cumulative variance %** | 24.12 | 44.45 | 60.31 | 70.49 | 79.23 |

**Supp Table 9: Principal component analysis of the measured morpho-physiological parameters and enzyme assays under ON conditions.** PRO: protein; CHLA: chlorophyll A; CHLB: chlorophyll B; TCHL: total chlorophyll; Caro: carotenoids; SL: shoot length; SFW: shoot fresh weight, SDW: shoot dry weight, RL: root length; RFW: root fresh weight
