## Supplementary.zip for "Unraveling the Interplay of Different Traits and Parameters Related to Nitrogen Use Efficiency in Wheat: Insights for Grain Yield Influence": Suppl Table 1.docx

| **GenID** | **Genotypes** | **GenID** | **Genotypes** | **GenID** | **Genotypes** | **GenID** | **Genotypes** |
| --- | --- | --- | --- | --- | --- | --- | --- |
| 1 | A 9-30-1 | 30 | GW 1245 | 59 | HD 2932 | 88 | HI 8704 |
| 2 | AKAW 4210 | 31 | GW 1280 | 60 | HD 2967 | 89 | HI 8713 |
| 3 | AKAW 4731 | 32 | GW 173 | 61 | HD 4502 | 90 | HI 8724 |
| 4 | AKW 1071 | 33 | GW 190 | 62 | HD 4672 | 91 | HI 8725 |
| 5 | AKW 381 | 34 | GW 273 | 63 | HD 4672 | 92 | HI 8727 |
| 6 | Altar 84 | 35 | GW 322 | 64 | HD 4676 | 93 | HI 8728 |
| 7 | AMRUT | 36 | GW 366 | 65 | HD 4709 | 94 | HI 8730 |
| 8 | Bijaga Red | 37 | GW 396 | 66 | HDR 77 | 95 | HI 8731 |
| 9 | Bijaga Yellow | 38 | HB 208 | 67 | HI 1077 | 96 | HI 977 |
| 10 | C 306 | 39 | HB 490 | 68 | HI 1500 | 97 | HINDI 62 |
| 11 | CHOTI LERMA | 40 | HD 1941 | 69 | HI 1531 | 98 | HP 1102 |
| 12 | DBP 01-09 | 41 | HD 1949 | 70 | HI 1539 | 99 | HP 1209 |
| 13 | DBP 01-11 | 42 | HD 1981 | 71 | HI 1544 | 100 | HP 1633 |
| 14 | DBW 14 | 43 | HD 1982 | 72 | HI 385 | 101 | HP 1731 |
| 15 | DBW 16 | 44 | HD 2009 | 73 | HI 7747 | 102 | HP 1744 |
| 16 | DBW 17 | 45 | HD 2135 | 74 | HI 784 | 103 | HP 1761 |
| 17 | DBW 39 | 46 | HD 2189 | 75 | HI 8381 | 104 | HPW 147 |
| 18 | DBW 71 | 47 | HD 2285 | 76 | HI 8498 | 105 | HPW 155 |
| 19 | DL 153-2 | 48 | HD 2329 | 77 | HI 8550 | 106 | HPW 226 |
| 20 | DL 784-3 | 49 | HD 2402 | 78 | HI 8591 | 107 | HPW 251 |
| 21 | DL 788-2 | 50 | HD 2643 | 79 | HI 8592 | 108 | HS 1097-17 |
| 22 | DL 803-3 | 51 | HD 2687 | 80 | HI 8627 | 109 | HS 1138-6-4 |
| 23 | DPW 621-50 | 52 | HD 2733 | 81 | HI 8638 | 110 | HS 240 |
| 24 | DWL 5023 | 53 | HD 2781 | 82 | HI 8645 | 111 | HS 277 |
| 25 | DWR 162 | 54 | HD 2824 | 83 | HI 8653 | 112 | HS 365 |
| 26 | DWR 195 | 55 | HD 2833 | 84 | HI 8663 | 113 | HS 375 |
| 27 | DWR-1006 | 56 | HD 2851 | 85 | HI 8671 | 114 | HUW 12 |
| 28 | DWR-2006 | 57 | HD 2864 | 86 | HI 8691 | 115 | HUW 206 |
| 29 | GW 1139 | 58 | HD 2888 | 87 | HI 8703 | 116 | HUW 234 |

| **GenID** | **Genotypes** | **GenID** | **Genotypes** | **GenID** | **Genotypes** |
| --- | --- | --- | --- | --- | --- |
| 117 | HUW 318 | 146 | KRL 237 | 175 | NIAW 34 |
| 118 | HUW 468 | 147 | LAL BAHADUR | 176 | NIDW 295 |
| 119 | HUW 55 | 148 | LERMA Rojo | 177 | NIPHAD 4 |
| 120 | HW 1085 | 149 | LOK 1 | 178 | NP 100 |
| 121 | HW 2004 | 150 | LOK 62 | 179 | NP 101 |
| 122 | HW 2045 | 151 | LOK 66 | 180 | NP 111 |
| 123 | HW 517 | 152 | MACS 1262 | 181 | NP 12 |
| 124 | HYB 633 | 153 | MACS 196 | 182 | NP 165 |
| 125 | HYB 65 | 154 | MACS 1967 | 183 | NP 4 |
| 126 | IWP 72 | 155 | MACS 2496 | 184 | NP 710 |
| 127 | J 405 | 156 | MACS 2846 | 185 | NP 715 |
| 128 | JNK-4W-184 | 157 | MACS 3125 | 186 | NP 718 |
| 129 | K 0307 | 158 | MACS 6145 | 187 | NP 737 |
| 130 | K 53 | 159 | MACS 6222 | 188 | NP 770 |
| 131 | K 68 | 160 | MOTIA | 189 | NP 771 |
| 132 | K 7410 | 161 | MP 1202 | 190 | NP 775 |
| 133 | K 7903 | 162 | MP 4010 | 191 | NP 792 |
| 134 | K 8020 | 163 | MPO 1106 | 192 | NP 799 |
| 135 | K 8027 | 164 | MPO 1215 | 193 | NP 809 |
| 136 | K 9006 | 165 | MPO 1236 | 194 | NP 818 |
| 137 | K 9107 | 166 | MPO 1255 | 195 | NP 824 |
| 138 | K 9533 | 167 | MPO 1259 | 196 | NP 825 |
| 139 | KALYANSONA | 168 | NARBADA 4 | 197 | NP 839 |
| 140 | KENPHAD 25 | 169 | NARMADA 112 | 198 | NP 846 |
| 141 | KENPHAD 39 | 170 | NI 5439 | 199 | NP 852 |
| 142 | KHARCHIA 65 | 171 | NI 5749 | 200 | NP 884 |
| 143 | KRL 1-4 | 172 | NI 917 | 201 | NP 890 |
| 144 | KRL 19 | 173 | NIAW 1549 | 202 | NW 1012 |
| 145 | KRL 229 | 174 | NIAW 301 | 203 | NW 1014 |

| **GenID** | **Genotypes** | **GenID** | **Genotypes** | **GenID** | **Genotypes** |
| --- | --- | --- | --- | --- | --- |
| 204 | NW 2036 | 233 | UAS 323 | 262 | VL 401 |
| 205 | PBW 175 | 234 | UAS 415 | 263 | VL 404 |
| 206 | PBW 222 | 235 | UAS 442 | 264 | VL 616 |
| 207 | PBW 226 | 236 | UASDW 30016 | 265 | VL 804 |
| 208 | PBW 343 | 237 | UASDW 30018 | 266 | VL 829 |
| 209 | PBW 373 | 238 | UASDW 30025 | 267 | VL 892 |
| 210 | PBW 396 | 239 | UASDW 30027 | 268 | VL 907 |
| 211 | PBW 443 | 240 | UASDW 30034 | 269 | WH 1021 |
| 212 | PBW 502 | 241 | UASDW 30037 | 270 | WH 1022 |
| 213 | PBW 550 | 242 | UASDW 30041 | 271 | WH 147 |
| 214 | PBW 590 | 243 | UASDW 30045 | 272 | WH 542 |
| 215 | PDW 215 | 244 | UASDW 30062 | 273 | WH 711 |
| 216 | PDW 233 | 245 | UASDW 30064 | 274 | WH 712 |
| 217 | PDW 291 | 246 | UASDW 30065 | 275 | WHD 896 |
| 218 | RAJ 1482 | 247 | UASDW 30071 | 276 | WL 410 |
| 219 | RAJ 1555 | 248 | UASDW 30075 | 277 | WL 711 |
| 220 | RAJ 1972 | 249 | UASDW 30076 | 278 | WR 544 |
| 221 | RAJ 3765 | 250 | UASDW 30078 |  |  |
| 222 | RAJ 3777 | 251 | UASDW 30079 |  |  |
| 223 | Raj 4037 | 252 | UASDW 30080 |  |  |
| 224 | RAJ 4083 | 253 | UASDW 30096 |  |  |
| 225 | RAJ 4248 | 254 | UASDW 30098 |  |  |
| 226 | RAJ 4294 | 255 | UP 115 |  |  |
| 227 | RW 346 | 256 | UP 2338 |  |  |
| 228 | SHARBATI SO | 257 | UP 2425 |  |  |
| 229 | SONALIKA | 258 | UP 2526 |  |  |
| 230 | SONORA 64 | 259 | UP 262 |  |  |
| 231 | Sujata (HI 617) | 260 | UP 301 |  |  |
| 232 | UAS 234 | 261 | UPD 93 |  |  |

**Supp Table 1: List of 278 selected wheat genotypes used in the study**
