## Supplementary.zip for "Unraveling the Interplay of Different Traits and Parameters Related to Nitrogen Use Efficiency in Wheat: Insights for Grain Yield Influence": Suppl Table 2.docx

| **Genotype** | **Year of release** | **Description (growing condition)** | **Yield (t ha^-1^) in AICRP trials** | | **Pedigree** |
| --- | --- | --- | --- | --- | --- |
|  |  |  | **Average** | **Potential** |  |
| NIPHAD-4 | 1942 | Resistant to alternaria blight | 1.2 | 2.0 | Motia(durum)/Khapli//NP4 (aestivum) |
| HUW-206 (MALAVIYA WHEAT 206) | 1985 | Resistance to all the  three rusts | 4.21 | 4.64 | KAVKAZ/BUHO//KAL YANSONA/BLUE BIRD |
| NP-852 | 1965 | Higher number of grains per spike | 3.50 | 4.00 | KF/2*NP 761 |
| C-306 | 1969 | Good for chapatti quality | 2.60 | 3.60 | RGN/CSK3 //2* C591/3/C217/N14 //C281 |
| HD-2967 | 2010 | Wider adaptability and resistance to yellow and brown rust | 5.0 | 6.6 | ALONDRA/CUCKOO//URES-81/HD-2160-M/HD-2278 |
| KENPHAD -39 | 1954-55 | Wider adaptability and resistance to rusts and susceptible to Alternaria blight | 2.5 | 3.5 | KENPHAD 25 SIB (Kenphad25 = K58F (L.1)/NI4) |
| RAJ-1482 | 1983 | Good for chapatti quality | 4.01 | 5.83 | NAPO-TOB 'S'/8156/KAL-BB |
| HD-2888 (PUSA WHEAT 107) | 2006 | High degree of resistance to brown rust, black rust and having high flour recovery | 2.25 | 3.83 | C306/T. *sphaerococcum*//HW 200 |
| GW-322 | 2002 | High degree of resistance to black rust, brown rust and tolerant to terminal heat | 4.4 | 6.6 | PBW-173/GW-196 |
| PBW-175 | 1989 | Resistance to yellow rust, brown rust and Karnal bunt | 2.80 | 4.81 | HD 2160 /WG 1025 |
| HS-365 | 1998 | Resistance to yellow rust | 1.82 | 2.68 | HS 207 /SONALIKA |
| K-7903 | 2001 | Tolerance to high  temperature | 2.5 | 3.5 | HD 1982/K816 |
| K-7410 | 1980 | Tolerance to drought and also suitable for alkaline soils | 3.23 | 4.64 | K 812 'S'/ KALYANSONA |
| RW-346 | 1989 | Stem rust resistant (carries Sr5, Sr8+ genes) | 2.5 | 3.5 | Janak/SA42 |
| WH-542 | 1992 | Higher content of zinc, copper and manganese | 4.8 | 6.1 | BJY/JUP//URES |
| WL-410 | 1978 | Resistance to yellow rust | 2.07 | 2.81 | SN63/4/36896//CJ 54/P4160E/3/HUA R/5/KAL |
| HP-1731 | 1995 | Resistance to brown  rust and tolerant to leaf  blight | 4.11 | 4.36 | LIRA //PARULA/  TONICH |

**Supp Table 2: Details of 17 selected wheat genotypes used in the study**
