## Supplementary.zip for "Unraveling the Interplay of Different Traits and Parameters Related to Nitrogen Use Efficiency in Wheat: Insights for Grain Yield Influence": Suppl Table 3.docx

| **S No** | **Gene** | **Primer Name** | **Primer sequence (5′ to 3′)** | **Length** | **Amplicon size** | **Tm** |
| --- | --- | --- | --- | --- | --- | --- |
| 1 | **Nitrite reductase** | NiR Frt | TGCCTCACCAAGAACAGC | 18 | 108 | 62.1 |
|  |  | NiR Rrt | CACCGCCTTCTTGTACACC | 19 |  | 62.4 |
| 2 | **Glutamine synthetase** | GS Frt | ATATGGTGAAGGGAACGAGC | 20 | 73 | 61.7 |
|  |  | GS Rrt | ACCCCATGAGAAGTCTGAAATG | 22 |  | 62.3 |
| 3 | **Glutamate synthase** | GOGAT Frt | AGGAGATTGAAGGATCACAAGAG | 23 | 130 | 61.8 |
|  |  | GOGAT Rrt | GCTTTGAAGTTGGAACGGTTG | 21 |  | 62.3 |
| 4 | **Glutamate dehydrogenase** | GDH Frt | GTGACGGTGAGCTACTTCG | 19 | 95 | 61.7 |
|  |  | GDH Rrt | CGGGTCATGTACGTCTTGAG | 20 |  | 61.8 |
| 5 | **Citrate synthase** | CS Frt | AGCACATTGGAAGGAGTCAC | 20 | 119 | 62.1 |
|  |  | CS Rrt | GTTAGATGCCACTCGTTCCC | 20 |  | 62.2 |
| 6 | **Nitrate reductase** | NR Frt | GAAAGGATACGCATACTCCGG | 21 | 119 | 62 |
|  |  | NR Rrt | CGTACTTGTTCGGCTTCTCC | 20 |  | 62.3 |
| 7 | **Isocitrate**  **dehydrogenase** | ICDH Frt | TTCCGTGTTCACCAGAAAGG | 20 | 140 | 62.3 |
|  |  | ICDH Rrt | GCTTCCTCAAGTTTCTGTGC | 20 |  | 61.3 |
| 8 | **Pyruvate kinase** | PK Frt | AAGCTGGTCGCCAAGTAC | 18 | 127 | 61.8 |
|  |  | PK Rrt | CTCTGTAGATCAGGCTGTGC | 20 |  | 61.7 |
| 9 | **Actin** | Actin Frt | GACCGTATGAGCAAGGAGATC | 21 | 122 | 61.8 |
|  |  | Actin Rrt | GTACTAAGGGAGGCAAGAATCG | 22 |  | 62.8 |

**Supp Table 3: Details of primers used for qPCR.**
